# Alterations of multiple alveolar macrophage states in chronic obstructive pulmonary disease

**DOI:** 10.1101/2020.05.28.121541

**Authors:** Kevin Baßler, Wataru Fujii, Theodore S. Kapellos, Arik Horne, Benedikt Reiz, Erika Dudkin, Malte Lücken, Nico Reusch, Collins Osei-Sarpong, Stefanie Warnat-Herresthal, Allon Wagner, Lorenzo Bonaguro, Patrick Günther, Carmen Pizarro, Tina Schreiber, Matthias Becker, Kristian Händler, Christian T. Wohnhaas, Florian Baumgartner, Meike Köhler, Heidi Theis, Michael Kraut, Marc H. Wadsworth, Travis K. Hughes, Humberto J. G. Ferreira, Jonas Schulte-Schrepping, Emily Hinkley, Ines H. Kaltheuner, Matthias Geyer, Christoph Thiele, Alex K. Shalek, Andreas Feißt, Daniel Thomas, Henning Dickten, Marc Beyer, Patrick Baum, Nir Yosef, Anna C. Aschenbrenner, Thomas Ulas, Jan Hasenauer, Fabian J. Theis, Dirk Skowasch, Joachim L. Schultze

## Abstract

Despite the epidemics of chronic obstructive pulmonary disease (COPD), the cellular and molecular mechanisms of this disease are far from being understood. Here, we characterize and classify the cellular composition within the alveolar space and peripheral blood of COPD patients and control donors using a clinically applicable single-cell RNA-seq technology corroborated by advanced computational approaches for: machine learning-based cell-type classification, identification of differentially expressed genes, prediction of metabolic changes, and modeling of cellular trajectories within a patient cohort. These high-resolution approaches revealed: massive transcriptional plasticity of macrophages in the alveolar space with increased levels of invading and proliferating cells, loss of MHC expression, reduced cellular motility, altered lipid metabolism, and a metabolic shift reminiscent of mitochondrial dysfunction in COPD patients. Collectively, single-cell omics of multi-tissue samples was used to build the first cellular and molecular framework for COPD pathophysiology as a prerequisite to develop molecular biomarkers and causal therapies against this deadly disease.

## Introduction

Worldwide, chronic obstructive pulmonary disease (COPD) is the third leading cause of death, with China and India accounting for more than 50% of all cases (Celli and Wedzicha, 2019; Rabe and Watz, 2017; Roth et al., 2018). In 2017, 3.2 million deaths due to COPD were estimated, a number that could increase to 4.4 million per year by 2040 (Celli and Wedzicha, 2019; Rabe and Watz, 2017). Due to smoking and increasing air pollution the current prevalence of 10.1% is estimated to further increase in the next decades (Celli and Wedzicha, 2019). Considering the enormous medical and financial burden of COPD (May and Li, 2015; Patel et al., 2018), there is a need to develop efficient biomarker-based diagnostics, and molecularly defined therapies. Current diagnosis and characterization of disease severity in COPD patients relies on clinical parameters with the baseline value for Forced Expiratory Volume in 1 second (FEV1) being the major parameter (Global Initiative for Chronic Obstructive Lung Disease (GOLD) Grade 1-4) (Celli and Wedzicha, 2019). However, it is now accepted that COPD is a heterogeneous disease manifesting as a clinical syndrome with structural pulmonary abnormalities, lung function impairment, chronic respiratory symptoms, or any combination of these (Celli and Agustí, 2018). Consequently, the pathogenesis of the disease is complex with numerous co-existing mechanisms (Agustí and Faner, 2018; Agustí and Hogg, 2019), including enhanced apoptosis (Voelkel et al., 2004), failure of lung tissue maintenance (Tuder et al., 2006), oxidative stress (Bowler et al., 2004; McGuinness and Sapey, 2017), protease/antiprotease imbalance (Abboud and Vimalanathan, 2008; Pandey et al., 2017), cellular senescence (Houssaini et al., 2018) and immunosenescence (Barnes, 2017). However, inflammation is the most prominent and important factor in the pathogenesis (Barnes et al., 2015). In comorbid patients, COPD is most likely the pulmonary component of chronic systemic inflammation (Fabbri and Rabe, 2007).

Lung inflammation in COPD is characterized by alterations in the number and function of immune cells, whereby macrophages and neutrophils gained much attention (Barnes, 2019). Elevated cell numbers in COPD have also been described for eosinophils (Berg and Wright, 2016; Saha and Brightling, 2006), CD8^+^ T cells (O’Shaughnessy et al., 1997) and innate lymphoid cells (ILCs) (De Grove et al., 2016). Despite the knowledge of elevated neutrophil counts in sputum and other biopsy specimens in COPD, therapeutic strategies targeting neutrophils have not been clinically effective. This suggests that a better understanding of the heterogeneous cellular immune response in COPD is still critical (Barnes, 2013). Interestingly, still little is known about cell-type heterogeneity, particularly during earlier disease grades. This is also true for the alveolar macrophage (AM) population, which is assumed to be highly heterogeneous (Kapellos et al., 2018), although a detailed characterization is still pending. AMs are considered to be one of the major orchestrators in COPD (Barnes, 2004), since, among other aspects, defective phagocytosis of bacteria is suspected to be associated with frequent exacerbations, a phase of acute worsening of the disease syndrome accompanied with hospitalization and increased mortality (Han et al., 2017; Hurst et al., 2010). Yet, the underlying molecular mechanisms are unresolved.

Heterogeneity of disease states and potential molecular mechanisms strongly argues for significant cellular heterogeneity, a research area which has gained much attention through the advent of single-cell genomics (SCG) technologies (Bassler et al., 2019; Chen et al., 2019; Giladi and Amit, 2018; Gomes et al., 2019; Linnarsson and Teichmann, 2016; Papalexi and Satija, 2018; Regev et al., 2017; Tanay and Regev, 2017; Wagner et al., 2016). Indeed, like for other organs, the cellular components of the human lung in health and disease have been the targets of an increasing number of studies. For example, SCG was used for the description of the parenchymal cell composition (Madissoon et al., 2020; Travaglini et al., 2019; Vieira Braga et al., 2019), the identification of novel cell types such as pulmonary ionocytes (Montoro et al., 2018; Plasschaert et al., 2018), the identification of ectopic and aberrant lung resident cell populations in idiopathic pulmonary fibrosis (Adams et al., 2019; Morse et al., 2019; Reyfman et al., 2019) or the investigation of the cellular contribution in lung cancer (Lambrechts et al., 2018; Lavin et al., 2017; Song et al., 2019; Zilionis et al., 2019). These studies in humans have been accompanied by SCG studies describing the cellular compositions in murine lung under homeostatic as well as stress conditions (Angelidis et al., 2019; Aran et al., 2019a; McQuattie-Pimentel et al., 2019; Schyns et al., 2019; Strunz et al., 2019), and during development (Cohen et al., 2018; Guo et al., 2019). Together, these studies illustrate the enormous breadth of SCG technologies to describe the cellular composition of the lung and identify deviations from homeostasis in diseased organ tissues.

Inspired by these promising results, we hypothesized that such high-resolution information will help to address many open questions in COPD biology, ranging from the involvement of different cellular compartments, to the role of immune cells for pathogenesis and progression, the distribution of cell-types within compartments, the molecular deviations involved in pathogenesis, but also clinical applicability of SCG technologies to define potential disease biomarkers in clinically relevant specimens such as bronchoalveolar lavage fluid (BALF) or peripheral blood. In this study, we focused our efforts on the immune compartment in BALF and blood in early-grade disease (GOLD grade 2). This early COPD grade has been recognized as potentially targetable grade to significantly reduce the incidence to develop more severe COPD. Collectively, we applied multi-color flow cytometry (MCFC), array-based single-cell transcriptomics derived from more than 160,000 cells, machine learning-assisted cell-type determination, and a novel differential expression (DE) analysis to define the molecular deviations in COPD patients at high resolution with a special emphasis on AMs. Several of the predicted deviations in lipid metabolism, antigen presentation, cellular motility, or oxidative stress response were experimentally validated, indicating the predictive value of SCG for a better understanding of COPD pathology.

## Results

### Single-cell transcriptomics of BALF from COPD patients

BALF is a clinically used biospecimen to assess the cellular compartment of the alveolar space in the context of lung diseases. Following previously defined criteria for using BALF in an explorative research setting (Meyer et al., 2012), we obtained high-quality BALF biopsy material and peripheral blood for comparative analyses (**Figure 1A**) from one COPD patient and four donors with chronic coughing but without any signs for pathophysiological alterations of the lung (hereafter referred to as ‘control’) (**Table S1**). Prior to applying single-cell RNA-sequencing (scRNA-seq) for high-resolution assessment of the cellular compartment in the alveolar space, we used MCFC based on 12 protein markers (**Figure S1A, Table S2)** to define the ground truth of the frequencies of major cell types to be expected in the AS (**Figure 1B**) in comparison to peripheral blood (**Figure S1B**). We used a strategy (**Figure S1A**) that is based on dimensionality reduction combined with dataset projection to generate an integrated UMAP graph to visualize the MCFC data. Most cells in the alveolar space were HLA-DR^+^ and highly autofluorescent, which - based on prior knowledge (Bharat et al., 2016; van Haarst et al., 1994; Vermaelen and Pauwels, 2004) - defined these cells as AMs (**Figure 1B, S1A**). These cells were absent in peripheral blood (**Figure S1B**). In addition to AMs, granulocytes (mainly neutrophils), monocytes, dendritic cells (DCs), NK cells, T cells, and few B cells were also detected in the alveolar space (**Figure 1B**). The cell population structure based on classical gating strategies of the MCFC data led to very similar results (*data not shown*). While MCFC was sufficient for describing major cell types, it lacked high resolution, which is necessary to identify cellular subsets or even functional states of individual subsets.

**Figure 1.**
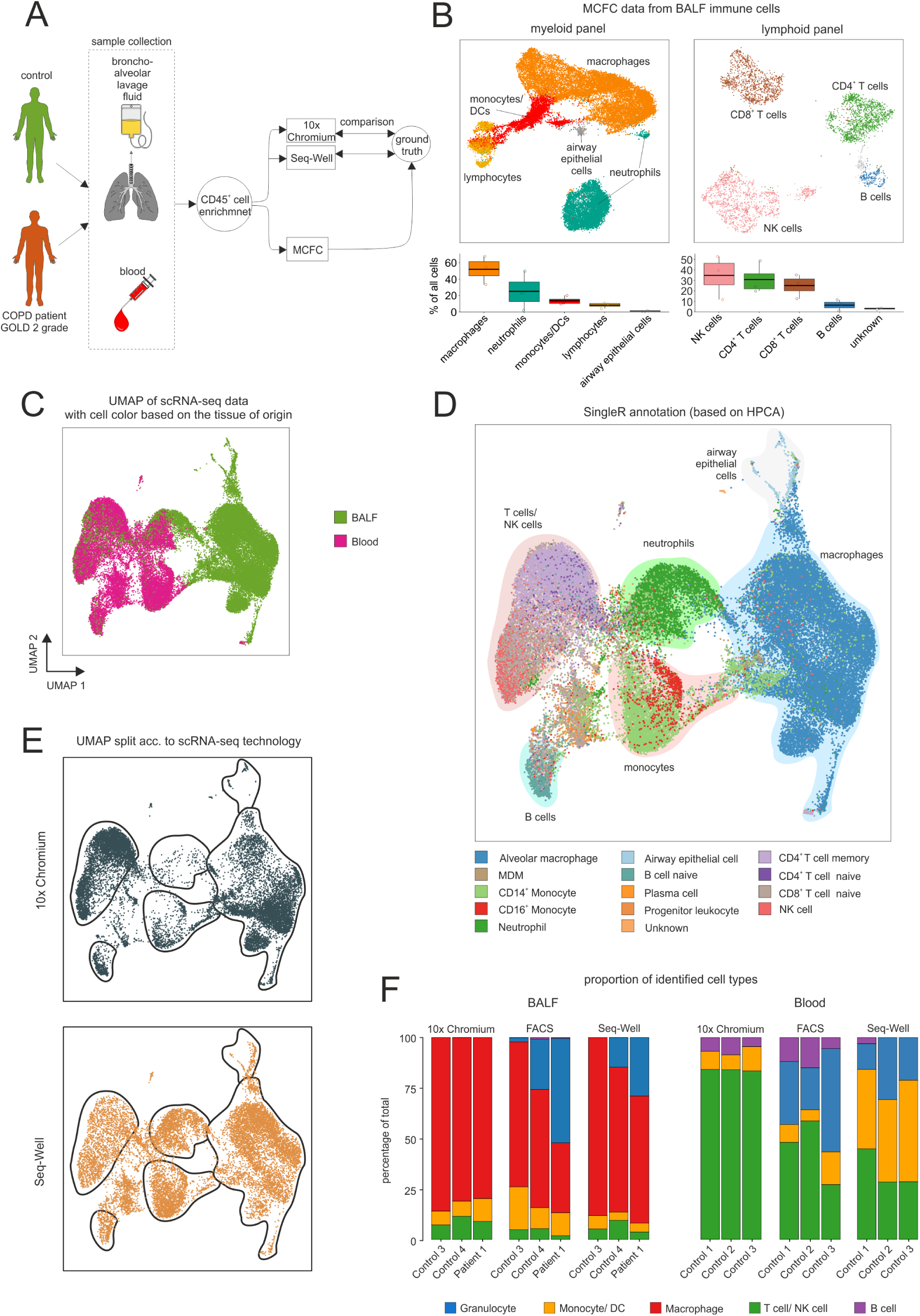
Evaluation of scRNA-seq data of BALF samples obtained from Seq-Well and 10x Chromium technology. **(A)** Schema describing the workflow of the comparison. **(B)** UMAP representation of MCFC data obtained from BALF immune cells of three donors. The relative proportion of identified cell types per donor displayed as boxplots. **(C)** UMAP representation of integrated blood and BALF data from different patients and the two scRNA-seq technologies. **(D)** Cell annotation based on the SingleR method with the HPCA dataset as reference (SingleR (HPCA)). Accumulation of cells with the same identity were highlighted with colored background and labeled accordingly. **(E)** UMAP representation of integrated data split by the two scRNA-seq technologies. The borders represent the areas of cell accumulation with the same identity as defined in (D)**. (F)** Stacked bar plots of the relative cell-type proportions for MCFC, which served as ground truth, and cell-type proportions, as predicted by SingleR (HPCA), of the two scRNA-seq technologies. BALF = bronchoalveolar lavage fluid; MCFC = multi-color flow cytometry; acc. = according; 10x Chrom. = 10x Chromium.

For clinical studies, the criteria to choose the most suitable scRNA-seq methodology include, among others: 1) capture of the major cell types, 2) appropriate scalability, 3) sufficient feature recognition, and 4) clinical applicability. Since some of the cell types that we identified in the alveolar space, such as granulocytes, do not withstand cryopreservation very well (Boonlayangoor et al., 1980), we chose to perform the study on freshly isolated biopsy material. To this end, we compared the most widely used droplet-based solution (Chromium from 10x Genomics Zheng et al., 2017) with a well-based method (Seq-Well (Gierahn et al., 2017)) using CD45^+^ immune cells from three BALF and three PB samples. In total, 48,531 cells (28,066 BALF cells and 20,465 blood cells) were assessed. Analyzing typical quality parameters in scRNA-seq data revealed slightly more uniquely aligned reads (**Figure S1C**) and a tendency to capture more cells (**Figure S1D**) for the well-based method, while the droplet-based method showed higher numbers of reads/cell (**Figure S1E**), transcripts/cell (**Figure S1F**), genes/cell (**Figure S1G**), and mitochondrial genes (**Figure S1H**). We integrated the scRNA-seq data from all samples by using a recently described anchoring strategy (Stuart et al., 2019) and visualized them via UMAP (Becht et al., 2019; McInnes et al., 2018) (**Figure 1C-E**). There were many immune cells found exclusively in BALF, whereby cells from the BALF were also found in areas defined mainly by blood cells (**Figure 1C**). We next annotated the cells using the SingleR method (Aran et al., 2019a) and the Human Primary Cell Atlas (HPCA) (Mabbott et al., 2013) as reference dataset, which identified most of the BALF-exclusive cells as macrophages with some contaminating CD45^+^ airway epithelial cells (**Figure 1D**). A similar result was achieved when using Blueprint (Stunnenberg et al., 2016) and ENCODE (Dunham et al., 2012) datasets as reference for SingleR. As a complementary approach, we aimed at annotating cells using gene signatures of an immune cell dataset (LM22) that have been described to robustly differentiate immune cell types (Newman et al., 2015). To enable annotation of single cells based on predefined gene signatures, we developed GenExPro, which uses a linear regression approach for cell annotation. Comparison of GenExpPro with the SingleR results showed that there is an general concordance between the different cell annotation methods, albeit not complete (**Figure S1J**). Overall, we identified all major immune cell types defined by MCFC (**Figure 1B, S1B**) also by scRNA-seq (**Figure 1D**). However, when determining the cell-type distribution for the droplet- and well-based scRNA-seq methods independently, it became obvious that the granulocytes (neutrophils, eosinophils) were almost completely lost in the droplet-based method (**Figure 1E, 1F, S1K**). While cell annotation methods showed slight differences (**Figure S1K**), all indicated the almost complete absence of granulocytes in the droplet-based method. Collectively, scRNA-seq by a well-based method combined with MCFC was the best choice for the determination and characterization of immune cells in BALF.

### Classification of immune cell types in the human alveolar space

For in-depth classification of cells within the alveolar space, we generated a second, larger Seq-Well-based scRNA-seq dataset from BALF samples derived from a cohort consisting of nine patients with early-grade COPD (GOLD 2) and six controls (**Table S2**). For cell annotation, we aimed to combine the results of GenExPro and SingleR to provide the best possible identification of cell types to answer clinically relevant questions, since consolidation allows the annotation not to be based on a single reference or method. For this reason, we developed a four-step approach comprising 1) a newly generated machine learning-based classifier, 2) cell clustering, followed by 3) a manual classifier-to-cluster comparison and 4) cluster-level marker gene analysis including cleanup (**Figure 2A**). The complete dataset (n = 60,925 cells) was visualized by UMAP representation revealing a highly heterogeneous structure within the integrated dataset (**Figure 2B**) that was not due to interindividual batch effects (**Figure S2A**).

**Figure 2.**
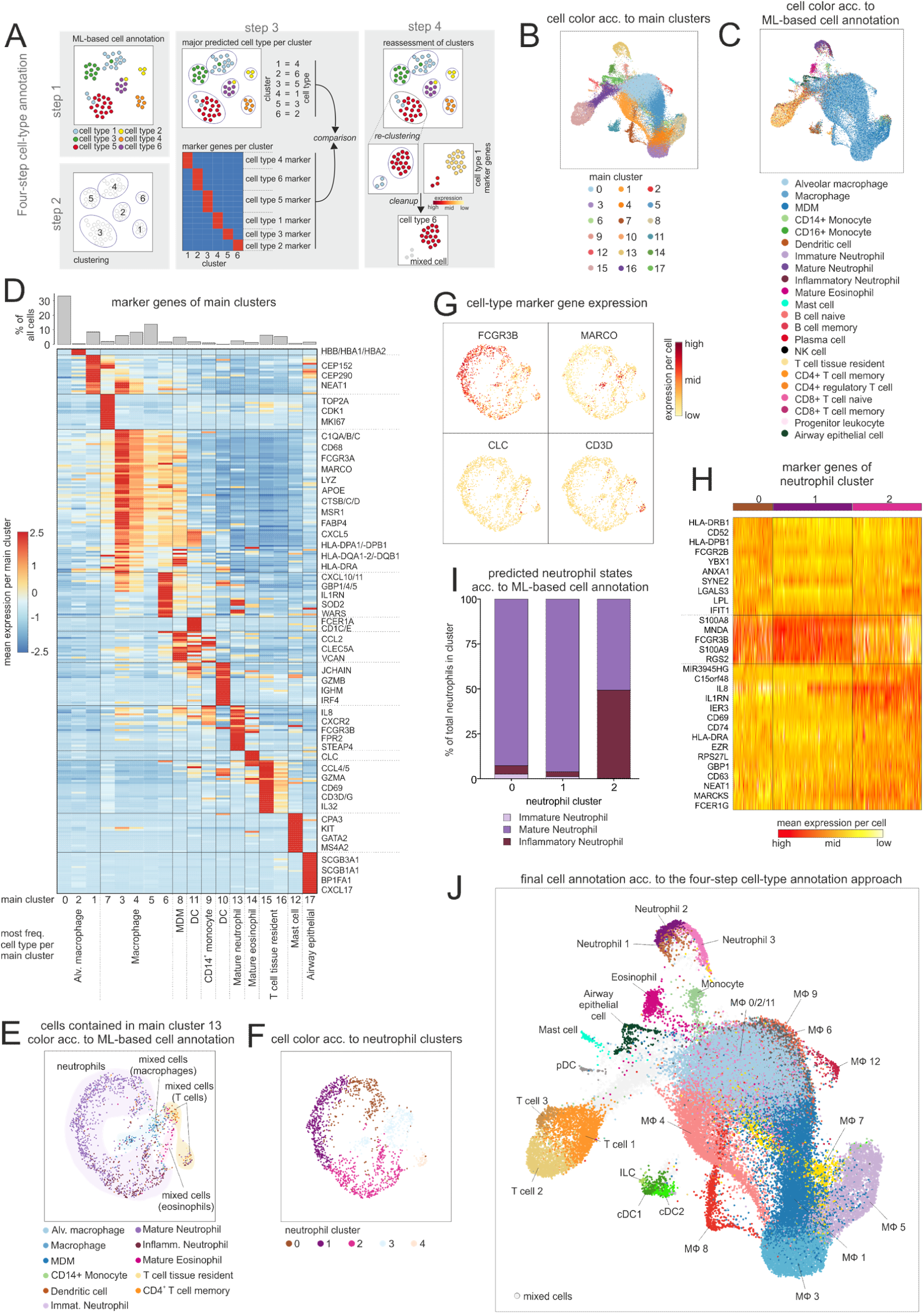
Machine learning-based cell type annotation of BALF immune cells. **(A)** Schematic workflow of the four-step annotation approach, including machine learning-based cell-type annotation, clustering, assignment and subsequent confirmation of a cluster to a cell type according to the machine learning-based cell-type annotation, and identification of ‘contaminating’ cells (referred to as ‘mixed cells’). **(B)** UMAP representation of integrated BALF data obtained from different COPD patients and control donors. Coloring according to identified main clusters. **(C)** UMAP representation of integrated BALF data with coloring according to machine learning-based cell-type annotation. **(D)** Heat map of the calculated marker genes per main cluster with a bar chart representation of the relative cell-type proportions at the top. The marker gene expression per cluster is represented as a z-transformed value (across all clusters). Rows of the heat map are sorted by hierarchical clustering. At the bottom of the plot, the main cell type according to the machine learning-based cell-type annotation is displayed, which is contained in the respective main cluster (acc. to Fig. S2F). **(E)** UMAP representation of cells contained in main cluster 13 (acc. to (B)). Coloring according to the machine learning-based cell-type annotation. Accumulation of cells with the same identity were highlighted with colored background and labeled accordingly. **(F)** UMAP representation and clustering of the cells contained in main cluster 13. **(G)** Feature plots showing the expression of cell type-specific markers (FCGR3B = neutrophils; MARCO = macrophages; CLC = eosinophils; CD3D = T cells). **(H)** Heat map of markers genes for neutrophil clusters 0-2 (acc. to (F)), which were predicted to contain mainly neutrophils according to the machine learning-based cell-type annotation (acc. to. (E)). Rows of the heat map are sorted by hierarchical clustering. **(I)** Bar chart showing the proportions of different neutrophil states in the neutrophil clusters 0-2 according to the machine learning-based cell-type annotation. **(J)** Final cell-type annotation of integrated BALF data according to the four-step annotation approach. BALF = bronchoalveolar lavage fluid; ML = machine learning; acc. = according; freq. = frequent; MDM = monocyte-derived macrophage; ILC = innate lymphoid cell; MΦ = macrophage.

As the first step, we applied our machine learning-based cell classifier that allowed us to aggregate knowledge over multiple cell-annotation algorithms in order to find consolidated cell labels (**Figure 2C and Figure S2B**). We tested the validity of this method using a benchmarking dataset generated by extracting cells with unequivocal expression of known cell type-specific markers from a reference scRNA-seq dataset (**Figure S2C+D**). Neither SingleR nor GenExPro alone were able to correctly annotate all cells within the benchmarking dataset, due to incomplete cell type-annotation within the reference (**Figure S1I**) used in these approaches. In contrast, our machine learning-based cell classifier was successful in consolidating the annotation results and thus resolving the different cell types (**Figure S2C**). Applying the machine learning-based cell annotation to the integrated BALF dataset revealed all major immune cell types and for some cell types a subset structure (**Figure 2C**).

The second step consisted of clustering of the data that resulted in 18 main clusters (**Figure 2B**), which agreed with the areas that were enriched for distinct cell types predicted by the classifier (**Figure 2C**). However, we also found some cells that were annotated, *e.g.* as neutrophils (**Figure S2E**), which scattered away from the other neutrophils.

Therefore, as the third step of the cell-type annotation procedure, we defined marker genes for each cluster (**Figure 2D**), defined the distribution of the classifier-defined major cell types within each cluster (**Figure S2F**) and re-assessed all cells within each of the 18 main clusters individually. As exemplified for main cluster 13 (neutrophils) (**Figure 2B**), we found striking concordance between the calculated marker genes and the predicted cell types that expressed the neutrophil markers *IL8, FCGR3B* and *CXCR2*, and most of the cells within the clusters were also annotated as neutrophils (**Figure 2D, Figure S2F**). Further investigation of main cluster 13 in the final step of the four-step cell-type annotation approach revealed that most of the cells were predicted by the classifier as neutrophils, and only a few cells were classified as T cells, macrophages or eosinophils (**Figure 2E**). Interestingly, the latter cells formed also distinct subclusters within cluster 13 (**Figure 2F**) and expressed known markers for the predicted cell types (macrophage = *MARCO*, eosinophils = *CLC*, T cells = *CD3D*) (**Figure 2G**), supporting the validity of the machine learning-based cell annotation. However, these ‘contaminating’ cells showed also expression of neutrophil markers (**Figure S2G**), indicating that these cells (hereafter referred to as ‘mixed cells’) might either represent putative cell doublets or low-quality cells and hence were excluded from further analysis. The remaining cells within this cluster, recognized as neutrophils, were analyzed more closely by calculating marker genes of the three identified subclusters (referred to as ‘neutrophil cluster’) (**Figure 2H**). Intriguingly, cells within neutrophil subcluster 2 expressed *CD63*, which has been linked to airway neutrophils in cystic fibrosis (Tirouvanziam et al., 2008). Moreover, the same cluster showed increased expression of activation markers, such as *CD69, FCER1G*, and *GBP1*. This is consistent with the labeling of almost half of the cells within this cluster as ‘inflammatory neutrophil’ by the machine learning-based cell annotation (**Figure 2I**).

We further underpinned the four-step cell-type annotation approach (**Figure 2A**) using main clusters 10 and 11 (**Figure 2B)** that were associated with dendritic cells (DCs). The large majority of the cells were classified as DCs (**Figure S2H**) but we also found an aggregation of ‘mixed cells’ (DC-cluster 0 = macrophage-associated, DC cluster 1 = monocyte-/MDC-associated) (**Figure S2H+I**). The major DC population formed distinct clusters (**Figure S2J**) representing cDC1, cDC2, and pDCs subpopulations, respectively, based on marker gene expressions (**Figure S2K**). In addition, we found one cluster (DC subcluster 5) that showed expression of *IL7R* and *CCL22* (**Figure S2K**) but not *CD3D* (**Figure S2I)**, which identifies these cells most likely as rare ILCs contaminating the DC cluster, further illustrating the power of our combined approach to even identify rare cell types. The application of this approach to all 18 major clusters in the integrated BALF dataset led to a detailed resolution of the immune landscape in the alveolar space (**Figure 2J**). In line with the FACS data of the alveolar space (**Figure 1B**), we identified macrophages as the major immune cell population (**Figure 2D, bar plot**). Together with monocytes, dendritic cells, mast cells, T cells, eosinophils, and neutrophils, they form the immune compartment in the alveolar space (**Figure 2J, Table S3**).

Collectively, our four-step approach starting with the machine learning-based classifier and ending with a rigid validation process for each cell in each major cluster 1) allowed us to remove any cell with ambiguous information content (approx. 9% were assigned as ‘mixed cells’ and thus removed from further analysis), and 2) led to the identification of cell states and rare cell types jointly across the 15 patients prior to using the dataset for addressing disease-related questions.

### Altered lipid metabolism in AMs of COPD patients

Our scRNA-seq analysis revealed the myeloid compartment with mononuclear cells (main clusters 0 – 9, **Figure 2B+D**) – comprising AMs - to be the largest cellular compartment in the alveolar space (**Table S3**). To elucidate functional heterogeneity and differences between control and COPD patients within these cells, we further investigated the compartment and identified 13 clusters (**Figure 3A**) of which one was clearly a monocyte cluster (corresponding to main cluster 9 (**Figure 2B**)) as indicated by the machine learning-based cell-type annotation (**Figure S3A).** Prior to defining cluster-specific functional changes between control and COPD, we searched for changes across the complete cluster space. For this purpose, we developed the so-called ‘GO-shuffling’ approach (**Figure S3B**). Briefly, this approach takes as input the average gene expression values per cell type of each patient and determines which functional gene sets, such as those based on gene ontology (GO) or pathway annotations, explain the strongest separation of COPD patients from controls in the Euclidean space. Enrichment analysis of functional terms within the upper percentile of the functional gene sets with the highest separation potential predicted by GO-shuffling per AM cluster showed that ‘metabolically’ associated terms mainly contributed to the separation of COPD patients from control donors. (**Figure 3B**). Interestingly, among other terms such as ‘activation’, ‘morphogenesis’, or ‘chemotaxis’, NOTCH signaling, was also found to be overrepresented (**Figure 3B**). A heat map representation of the expressed genes contained in the NOTCH gene sets confirmed that, for example, metalloproteases of the ADAM family (*ADAM17* and *ADAM9*) and the components of the γ-secretase complex (*APH1B, APH1A, PSEN1, PSENEN*, and *NCSTN*) are able to separate COPD patients from control donors (**Figure S3C**). To further examine potential COPD-dependent changes in metabolism, we applied the Compass algorithm (Wagner et al., 2020; Wang et al., 2020) to comprehensively model the metabolic differences between COPD and control AMs (**Figure 3C**). The largest differences were found in amino acid and lipid metabolism (**Figure 3C, pie chart**), with an overall higher predicted metabolic activity in COPD samples (**Figure 3C, heat map**). Among the differential lipid-associated metabolites and reactions, phosphorylation of inositol was most prominent, but we also found altered metabolites and reactions indicating increased transport (monoacylglycerol), synthesis (phospholipids and cholesterol) and degradation (β-oxidation) of lipids in COPD AMs.

**Figure 3.**
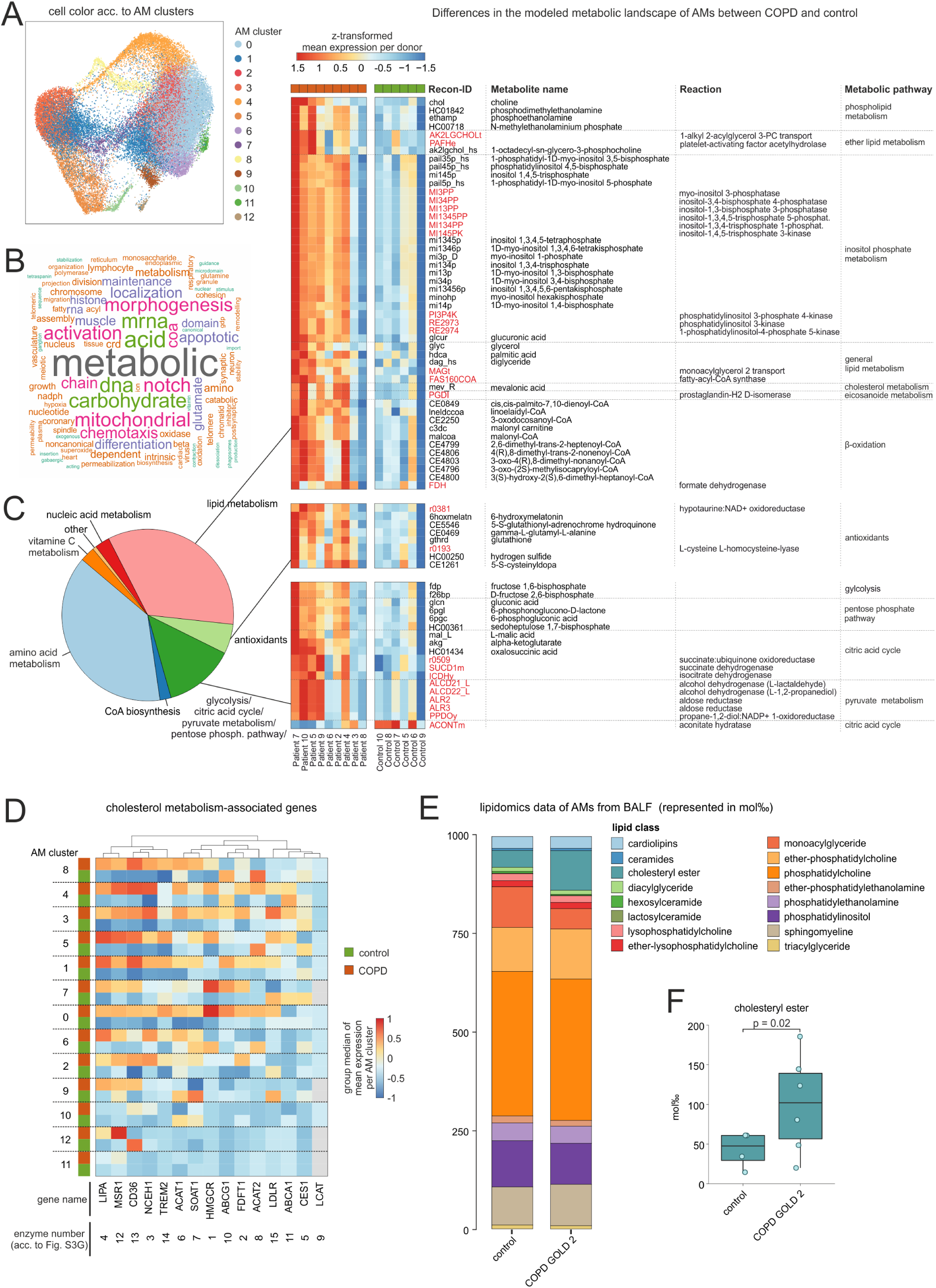
Modeling of the metabolic landscape in AMs. **(A)** UMAP representation and clustering of cells contained in main clusters 0-9, which are annotated as monocytes or macrophages (acc. to Fig. 2B+C). **(B)** Word cloud of the most common words in the top predicted terms of the GO-shuffling approach across all macrophage clusters. **(C)** Compass results of the modelled metabolic landscape in AMs. The Venn diagram summarizes and categorizes the predicted metabolites and pathways that are significantly different between COPD and control. Heat map shows the predicted pathways and metabolites associated with lipid metabolism, antioxidants and energy metabolism. Recon2-ID (Thiele et al., 2013) of metabolites is shown in black and reactions in red. Columns and rows of the heat map are sorted by hierarchical clustering. **(D)** Heat map representation of cholesterol metabolism-associated genes (Fig. S3G) across the identified macrophage/monocyte clusters (referred to as ‘AM clusters’) (acc. to (A)). Depicted is the group median (group = COPD or control) of the z-transformed mean expression data per donor and AM cluster across all AM clusters. Columns and rows of the heat map are sorted by hierarchical clustering. **(E)** Stacked bar displaying the mean proportions (represented in mol permille) of lipid classes obtained by lipidomics analysis (control n = 4, COPD n = 6). **(F)** Box plot of cholesterylester proportions (acc. to (E)) with the representation of individual donors. BALF = bronchoalveolar lavage fluid; acc. = according; AM = alveolar macrophage.

To evaluate the Compass-based prediction, we extracted the part of the phospholipid metabolism from the KEGG database, which, according to the Compass results, presented higher predicted metabolic activity in COPD than in the controls (**Figure S3D**). In accordance with Compass, we found that more cells express the genes encoding the enzymes involved in the predicted phospholipid pathway in COPD (**Figure S3E**). In addition, we validated the predicted elevation in lipid metabolism by visualizing the expression of genes found in lipid-associated gene sets contained in the upper percentile of the functional gene sets with the highest separation potential according to GO-shuffling (**Figure S3F**). Among these genes, we found several receptors for cholesterol uptake (*CD36, LDLR, MSR1*, and *TREM2*) and genes of cholesterol storage mediated by cholesteryl ester synthesis (*ACAT1/2* and *SOAT1*), but also genes encoding cholesteryl ester hydrolases (*LIPA* and *NCEH1*) (**Figure S3F**). To gain a better understanding of how cholesterol metabolism-associated genes (**Figure S3G**) are regulated in the identified clusters of the myeloid compartment (**Figure 3A**), we determined and plotted the median gene expression of COPD patients and control donors per cluster (**Figure 3D**). Interestingly, the smaller clusters (cluster 9 - 12) showed almost no difference between COPD and control, while the majority of genes in the larger clusters (cluster 0 - 8) showed a marked difference, with mostly higher expression in COPD. However, some genes were regulated differently across the clusters, such as the cholesterol acyltransferase *ACAT2*, which was downregulated in COPD in cluster 5, 6 and 8, but either upregulated or not deregulated in the remaining clusters.

Next, we validated the *in-silico* prediction of altered lipid metabolism in COPD by performing lipidomics analyses of 229 lipid species in AMs obtained either from COPD GOLD 2 patients or from control donors. We observed the greatest difference in the lipid classes for cholesteryl ester, which was higher in COPD AMs than in controls (**Figure 3E-F**). Based on this finding, we postulate that the cells within the myeloid cell compartment in COPD patients show a pulmonary foam cell-like response, which has also been reported for other lung diseases, such as lipoid pneumonia (Collins et al., 1995) or vaping-related lung injury (Maddock et al., 2019). This functional response is characterized by the cells being predominantly cholesterol-laden. In pulmonary alveolar proteinosis, increased cholesterol/lipid accumulation within AMs might be mediated by defective GM-CSF signaling and, as a consequence, reduced *PPARG* and cholesterol transporter (*ABCG1*) expression (De Aguiar Vallim et al., 2017; Sallese et al., 2017; Trapnell et al., 2019). This mechanism is unlikely in COPD since we do not observe clear downregulation of either *PPARG* (**Figure S3F)** or *ABCG1* (**Figure 3D**).

Taken together, we found metabolic changes in the AMs of COPD patients by scRNA-seq and validated this observation by lipidomics analysis, which showed an accumulation of cholesteryl ester in COPD GOLD 2.

### COPD leads to major changes in most AM states

To further define the functionalities of the myeloid cell compartment with mononuclear cells in the alveolar space, we calculated marker genes for the clusters defined in **Figure 3A**. Except for cluster 10, which was constituted by monocytes, this cell compartment clearly belonged to the macrophage cell lineage as defined by the expression of typical signature genes (*MSR1, MRC1, MARCO).* Yet, these cells displayed a remarkable transcriptional plasticity, which was reflected in the number of the identified clusters (**Figure 4A+B**). In addition to the macrophage signature, cluster 8 was also characterized by proliferation-associated genes (*MKI67, TOP2A*, and *NUSAP1*) as well as increased expression of histones (*HIST1H4C* and *HIST1H1D*). Furthermore, most of the cells within this cluster were assigned to the G2/M cell cycle phase (**Figure S4A**), strongly supporting that this cluster represented proliferating AMs. Clusters 9 and 6 were highly enriched for the expression of MHC class II molecules, namely HLA-DQ and HLA-DR respectively, while cluster 12 carried hemoglobin genes (*HBA2, HBA1*, and *HBB*) either due to engulfed erythrocytes or induction of hemoglobin genes in macrophages (Liu et al., 1999). We also examined whether the AM clusters were formed uniformly by all donors or whether a cluster was defined by the overrepresentation of an individual donor (**Figure S4B**). Only for cluster 12 (MΦ/erythrocyte), 2 and 11, we found a donor effect, with the latter being characterized by the expression of the T cell-associated genes *CD2* and *CCL5*, which led us to label this cluster as ‘ILC-like’ macrophages. Interestingly, cluster 5 exhibited relatively strong expression of the monocyte-associated genes *VCAN* and *S100A8* together with the monocyte attractant *CCL2* and the late monocyte-to-macrophage differentiation marker *CHIT1* and was therefore defined as ‘mono-like’ macrophages. This cluster also shared some markers with cluster 7, which was additionally high in interferon-response genes (*IFIT1* and *IFIT2*), and cluster 3 that showed increased expression of complement components (*C1QA-C*) and alpha-1-antitrypsin (*SERPINA1*). As an alternative approach, we predicted the functions of each cluster by gene set variation analysis (GSVA) (Hänzelmann et al., 2013a) (**Figure S4C**) and visualized the identified terms in an UpSet plot (Conway et al., 2017) (**Figure S4D**) illustrating shared but also cluster-specific functions. Among the shared terms, we found enrichment of ‘antigen presentation’, ‘endocytosis’, ‘oxidative phosphorylation’ and ‘β-oxidation’, which reflect some of the basic cellular processes of macrophages in the alveolar space. Intriguingly, cluster 4 revealed a specific enrichment of the mTOR signaling pathway, which was recently described to be associated with cellular senescence in non-immune cells from the lung (Barnes, 2017; Houssaini et al., 2018). To determine whether cluster 4 cells are indeed in a senescent state, we performed enrichment analysis of gene sets associated with cellular senescence (**Figure S4E**). We found an enrichment of cell aging and mitochondrial genes in cluster 4. In addition, the specific downregulation of genes described as downregulated in aged immune cells (IMM-AGE signature (Alpert et al., 2019)), supported the characterization of cluster 4 as senescent AMs (**Figure 4A+B**).

**Figure 4.**
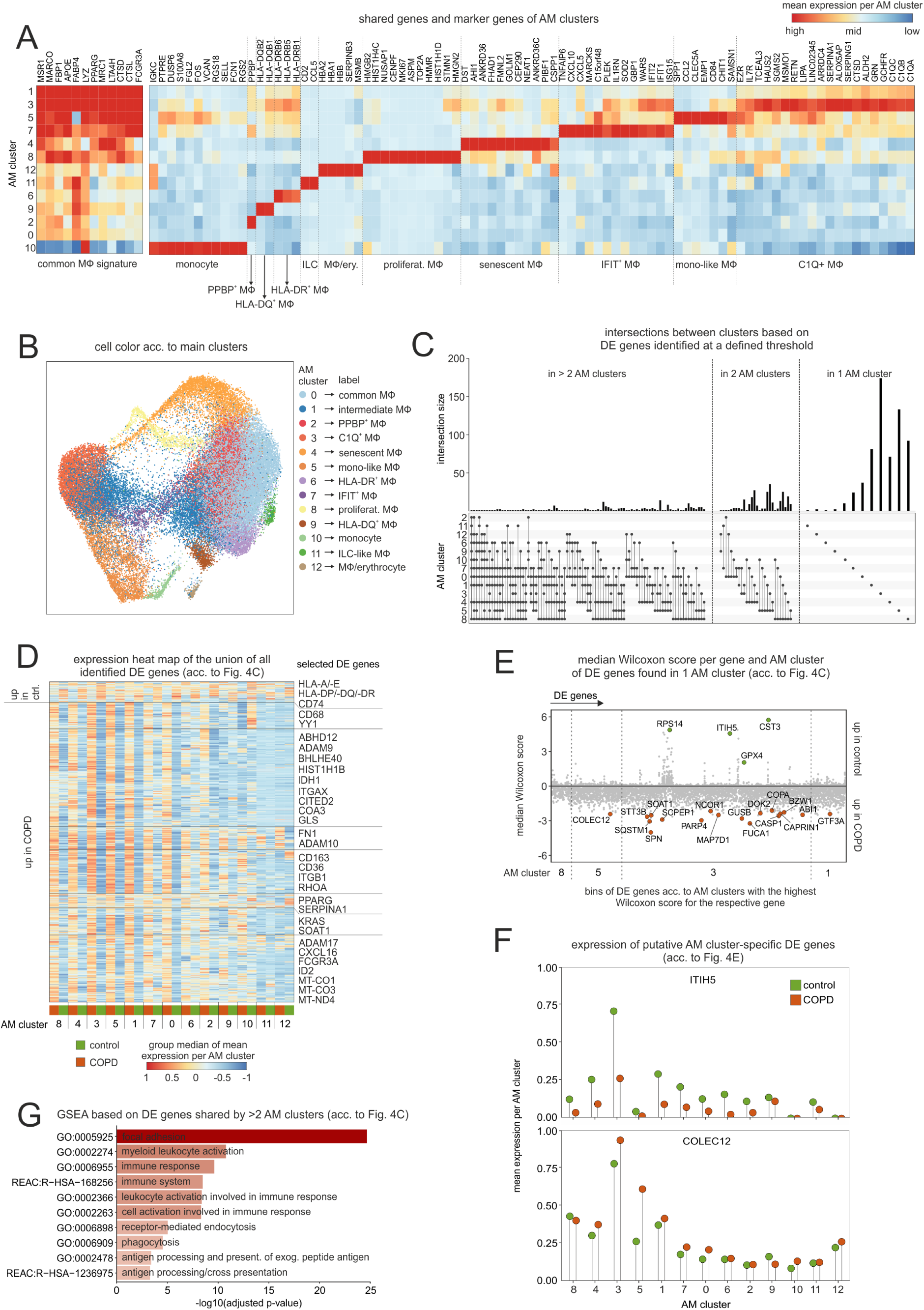
DE gene analysis of identified AM subtypes. (**A**) Heat map of calculated marker genes per macrophage/monocyte cluster (referred to as ‘AM cluster’; acc. to Fig. 3A). The marker gene expression per AM cluster is represented as a z-transformed value (across all AM clusters). On the left side of the heat map, conserved macrophage markers are depicted. Columns and rows of the heat map are sorted by hierarchical clustering. **(B)** UMAP representation of integrated macrophages and monocytes with coloring according to identified clusters. The clusters are labeled based on information from marker genes (acc. to (A)). **(C)** UpSet plot of calculated DE genes across clusters. DE genes found in the same clusters are binned and the size of the bins is represented as a bar chart. At the bottom, dots indicate which clusters contained and shared these DE genes. **(D)** Heat map representation of the union of all DE genes found in the AM cluster. Depicted is the group median (group = COPD or control) of the z-transformed mean expression data per donor and AM cluster across all AM clusters, and the names of some selected DE genes are shown on the right side of the plot. Columns and rows of the heat map are sorted by hierarchical clustering. **(E)** Dot plot for the assessment of the AM cluster specificity of DE genes. DE genes found in only one cluster according to UpSet plot in (C) are depicted on the x-axis and the respective Wilcoxon scores for each AM subtype on the y-axis. Dots of DE genes are highlighted (green = up in control; orange = up in COPD) and the respective gene name is shown if the p-value for the significance test (acc. to Fig. S4F) of the DE gene in the respective AM cluster is < 0.01 and its Wilcoxon score has a difference of ≤ 2 to the median Wilcoxon score of the remaining AM. In addition, DE genes with the highest Wilcoxon score in the same AM cluster are binned. The corresponding AM cluster number is shown at the bottom of the plot and dashed lines separate the bins. **(F)** Lollipop plot of two selected DE genes from (E)**. (G)** Selected functional gene sets from GSEA based on DE genes that reach the defined significance cutoffs for more than two AM subtypes (acc. to (C)). acc. = according; AM = alveolar macrophage; ILC = innate lymphoid cell; MΦ = macrophage; mono = monocyte; DE = differentially expressed; GSEA = gene set enrichment analysis.

Next, we examined each identified AM cluster for statistically significant transcriptional differences between COPD and controls. The present dataset represents an experimental setup that is currently emerging in the field of scRNA-seq, particularly in studies aimed at answering clinically relevant questions, namely replicated multi-condition datasets. To meet the needs of analyzing such scRNA-seq data, we developed a DE analysis approach based on the Wilcoxon rank sum test between COPD and control cells in combination with permutation tests that also considered possible individual donor effects (**Figure S4F**). Visualization of the identified DE genes per cluster in an UpSet plot (Conway et al., 2017) indicated that the majority of the observed transcriptional differences are cluster-specific (**Figure 4C**). However, further analysis revealed that the putative cluster-specific DE genes also exhibited differences with similar directions between COPD and controls in other clusters (**Figure 4D**). Similarly, plotting of the calculated Wilcoxon scores per gene also showed that the direction of the scores was the same across the clusters, mainly towards increased expression in COPD (**Figure 4E**). To test the validity of our DE analysis approach, we selected two genes (*ITIH5, COLEC12*) with a remarkably high Wilcoxon score in one cluster compared to the others and visualized their mean expression (**Figure 4F**). The difference between COPD and the control of the selected genes was particularly high in the respective clusters, which were also predicted by the DE analysis approach (**Figure 4E)**. Furthermore, the plot showed again that the expression trend was conserved across several clusters. The DE analysis revealed that most of the identified differences in gene expression between COPD and control cells were most pronounced in certain AM clusters, although the trends in expression differences were conserved across several AM clusters.

Among the DE genes, we identified NOTCH-associated genes such as *YY1* and the metalloendopeptidases *ADAM9, ADAM10* and *ADAM17* (**Figure 4D).** In accordance with the Compass analysis, lipid metabolism-associated genes (*e.g. CD36, COLEC12, SOAT1*, and *PPARG*) were upregulated in COPD, which was also true for genes associated with oxidative phosphorylation (*e.g. COA13, MT-CO2, MT-ND2*, and *MT-ATP6*) (**Figure 4D-E**). Moreover, we found an increased expression of the surface molecule *CD163* in AMs of COPD patients, which has been described previously for this disease and cell type by immunohistochemistry (Kaku et al., 2014; Kunz et al., 2011). To put the identified DE genes into a functional context, we performed gene set enrichment analysis (GSEA), which revealed an enrichment of terms associated with focal adhesion and immune response such as antigen processing and presentation (**Figure 4G**).

In conclusion, we were able to characterize the AM clusters, revealing a high transcriptional plasticity of this cell population. The characterized AM clusters are hereafter referred to as AM states. Furthermore, DE analysis showed that the transcriptional differences are mainly attributable to increased expression in COPD. In addition, we found that for the majority of the identified DE genes, the direction of the difference between COPD and control AMs was shared across several clusters, indicating a global and similar impact of the disease on different AM states in the alveolar space. Finally, further analysis revealed that some of the identified DE genes are associated with focal adhesion and antigen presentation.

### Reduced motility and MHC expression together with mitochondrial dysfunction of AMs in COPD

DE analysis across patient groups in the myeloid compartment of the alveolar space suggested alterations in focal adhesion and antigen presentation in COPD (**Figure 4D+G**). The DE genes associated with focal adhesion that were elevated in COPD, included cell adhesion molecules (*e.g. ITGB1* or *ALCAM*) and cytoskeleton organizing molecules (*e.g. PARVG* or *ARPC2/5*) (**Figure 5A**), pointing towards either increased motility, or increased local adhesion strength of these cells. At the same time, we observed considerable increases of chemokines and other soluble proteins in BALF in COPD patients including CCL3 known to be involved in motility and migration of macrophages (**Figure 5B**) (Opalek et al., 2007). As indicated by scRNA-seq, CCL3 can be derived from numerous cell types in the alveolar space of COPD patients, especially from neutrophils, but also from T cells, IFIT^+^ and mono-like macrophages (**Figures S5A**). However, despite the increase of CCL3 in BALF, the intrinsic property of AMs from COPD patients was an overall reduced migratory response towards CCL3 (**Figure 5C**), most likely due to increased adhesive properties.

**Figure 5.**
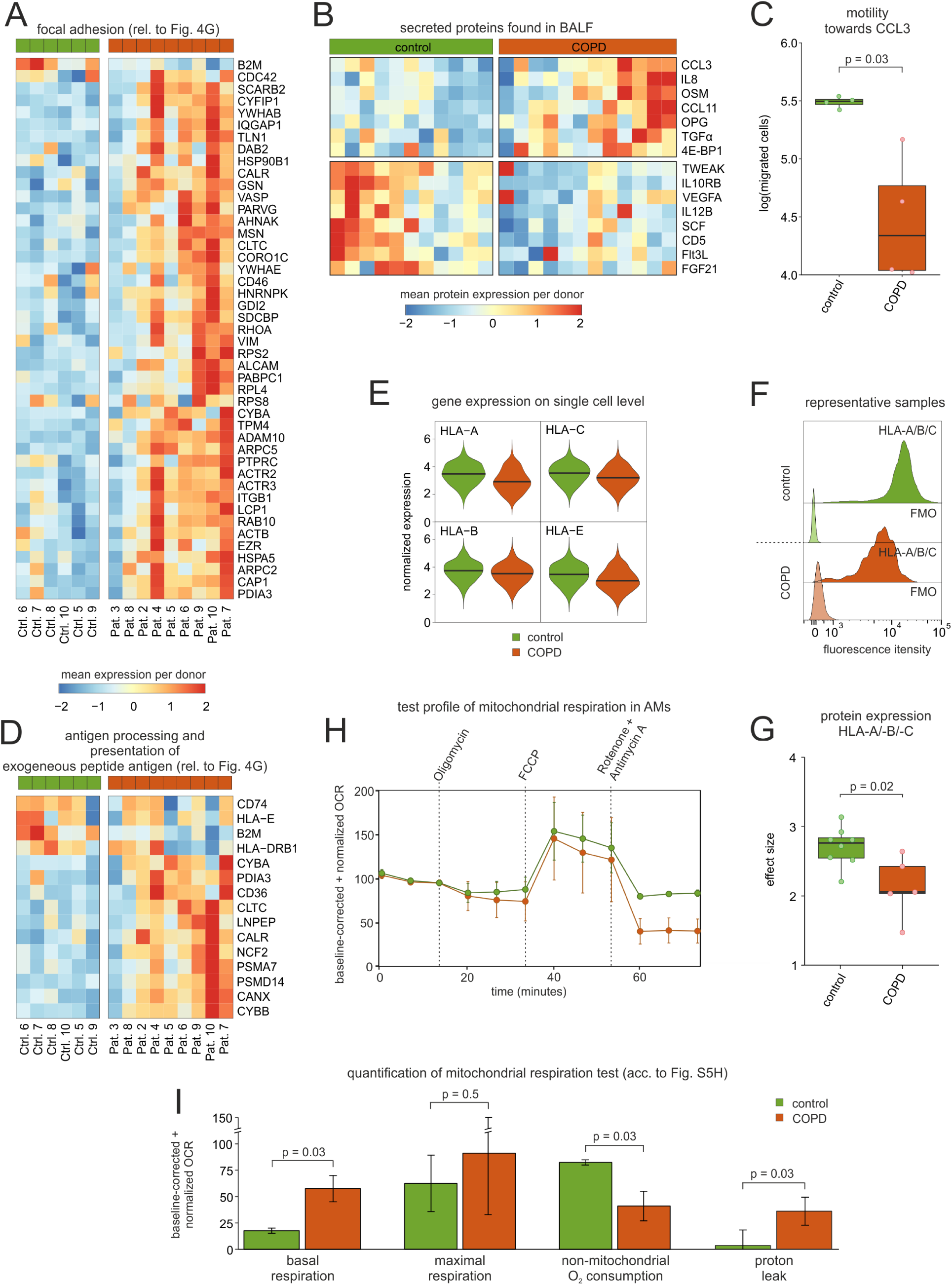
Assessment of antigen presentation, chemotaxis and mitochondrial function in AMs. **(A)** Heat map of DE genes, which according to GSEA are enriched in the GO term associated with focal adhesion (acc. to Fig. 3G). The mean gene expression per donor is represented as a z-transformed value (across all donors). Rows of the heat map are sorted by hierarchical clustering. **(B)** Heat map representation of proteins detected in BALF with a p-value < 0.1 according to the Wilcoxon rank sum test between COPD patients and control donors (control n = 11, COPD n = 12). The mean protein expression (identified by Olink Proteomics) per donor is represented as a z-transformed value (across all donors). Columns of the heat map are sorted by hierarchical clustering. **(C)** Quantification of the migratory capability of AMs towards CCL3 displayed in a box plot with the representation of individual donors (control n = 4, COPD n = 4). **(D)** Heat map of DE genes, which according to GSEA are enriched in the GO term ‘antigen processing and presentation of exogenous peptide antigen’ (acc. to Fig. 3G). The mean gene expression per donor is represented as a z-transformed value (across all donors). Rows of the heat map are sorted by hierarchical clustering. **(E)** Violin plot of the HLA-A expression in AMs based on scRNA-seq data. The plot shows the expression across the donors, whereby the donors were downsampled to the same number of cells, followed by downsampling to the same number of cells between COPD and control. The plot displays cells with an expression > 0. **(F)** Fluorescence intensity histograms showing representative samples of flow cytometric analysis of HLA-A/-B/-C expression on the cell surface of isolated AMs (FMO = fluorescence minus one). **(G)** Box plots of the calculated effect sizes of HLA-A/-B/-C expression in COPD and control with the representation of individual donors (control n = 8, COPD n = 5). **(H)** Evaluation of mitochondrial function via the time-dependent course of the oxygen consumption rate (OCR) in AMs, using baseline-corrected values. Error bars indicate the standard deviations derived from the measurements of several donors (control n = 2, COPD n = 3). Dashed arrows represent the injection of various compounds (shown at the top of the plot) used to assess different aspects of mitochondrial function (acc. to Fig. S5G). **(I)** Bar plots showing quantifications of different aspects of mitochondrial function inferred from the OCR measurement in (G) (acc. to Fig. S5G). BALF = bronchoalveolar lavage fluid; AM = alveolar macrophage; rel. = related

Next, we plotted DE genes associated with antigen presentation according to GSEA (**Figure 4G, Figure 5D**). We observed lower expression of the human leukocyte antigen (HLA) genes *HLA-E* and *HLA-DRB1*, and the HLA-associated genes *B2M* and *CD74* in COPD patients. To assess whether the altered expression could also be observed in other HLA genes, we comprehensively evaluated the expression profiles of MHC II and I cell surface receptors of AMs. Among the top expressed HLA genes, we found further genes of the MHC II class (*HLA-DRA, HLA-DRB1, HLA-DRB5, HLA-DPA1, HLA-DPB1*, and *HLA-DQB1*) together with MHC I-encoding genes (*HLA-A, HLA-B, HLA-C*, and *HLA-E*) (**Figure S5B**). Intriguingly, when plotting the expression of these genes for COPD and control samples separately, we found that the HLA genes were consistently downregulated in the COPD samples, with the only exception of HLA-DRA that showed no difference (**Figure 5E and Figure S5C**). We therefore isolated AMs from additional patients and measured protein levels of MHC class I (HLA-A/-B/-C) (**Figure 5F, 5G**), class II (HLA-DR) (**Figure S5D, S5E**), and CD74 (**Figure S5F**). Clearly, MHC class I and CD74 were significantly reduced on AMs of COPD patients, while MHC class II molecules showed no clear difference.

Finally, AMs of COPD patients were characterized by increased expression of mitochondrial genes (**Figure 4D**), which might represent a cellular adaptation to elevated metabolic activity as indicated by the Compass analysis (**Figure 3C**). To test this hypothesis, we isolated AMs from BALF of additional patients and controls and performed the MitoStress assays by the Seahorse technology (**Figure 5H and S5G**). Indeed, we identified increased baseline respiration in AMs of COPD patients (**Figure 5I**), which reflects an elevated energy demand. We also found a significant increase in proton leakage in AMs of COPD patients, which is indicative for increased ROS production in COPD (Boukhenouna et al., 2018; Cheng et al., 2017; McGuinness and Sapey, 2017) and mitochondrial dysfunction (Eapen et al., 2019; Hoffmann et al., 2019; Ng Kee Kwong et al., 2017).

Collectively, these validation studies verified the scRNA-seq results, which revealed the complex pathophysiological processes operative in COPD.

### A complex cell-to-cell communication network coordinates the regulation of DE genes in AMs

As a next step, we were interested in understanding the regulation of the identified DE genes and thus in defining intra- and inter-cellular signaling cascades. First, we focused on the myeloid compartment, particularly those AM states with a minimum of 30 DE genes between COPD and control and predicted potential upstream transcriptional regulators of the DE genes (**Table S3**). Representation of the predicted transcriptional regulators in an UpSet plot revealed that *YY1* was the only transcription factor (TF) shared among all AM states included in the analysis (**Figure 6A**), known to be an important modulator of NOTCH signaling (Liao et al., 2007; Yeh et al., 2003). Elevated NOTCH signaling in COPD was further supported by the identification of the TFs *HES1* and *HEY1* with co-regulation being present in mono-like macrophages and C1Q^+^ macrophages. In addition to NOTCH signaling, other predicted signaling cascades included WNT signaling (*e.g. TCF3/4, MYC* and *NFATC1/3*), TNF/ NF-κB signaling (*e.g. CEBPB* and *REL*), TGFβ signaling (*e.g. TFE3* and *MYOD1*), and TFs involved in the regulation of the circadian rhythm (*e.g. BHLHE40/41, CLOCK* and *TIMELESS*). Simultaneous elevation of these interconnected signaling pathways was further supported by the identification of TFs such as *MYC*, central to several pathways such as WNT, NOTCH, and TGFβ signaling. However, it also needs to be emphasized that other signaling cascades seemed to be much more specific for individual macrophage states, for example, the majority of interferon regulatory factors (IRFs) were only predicted for IFIT^+^ and proliferating macrophages.

**Figure 6.**
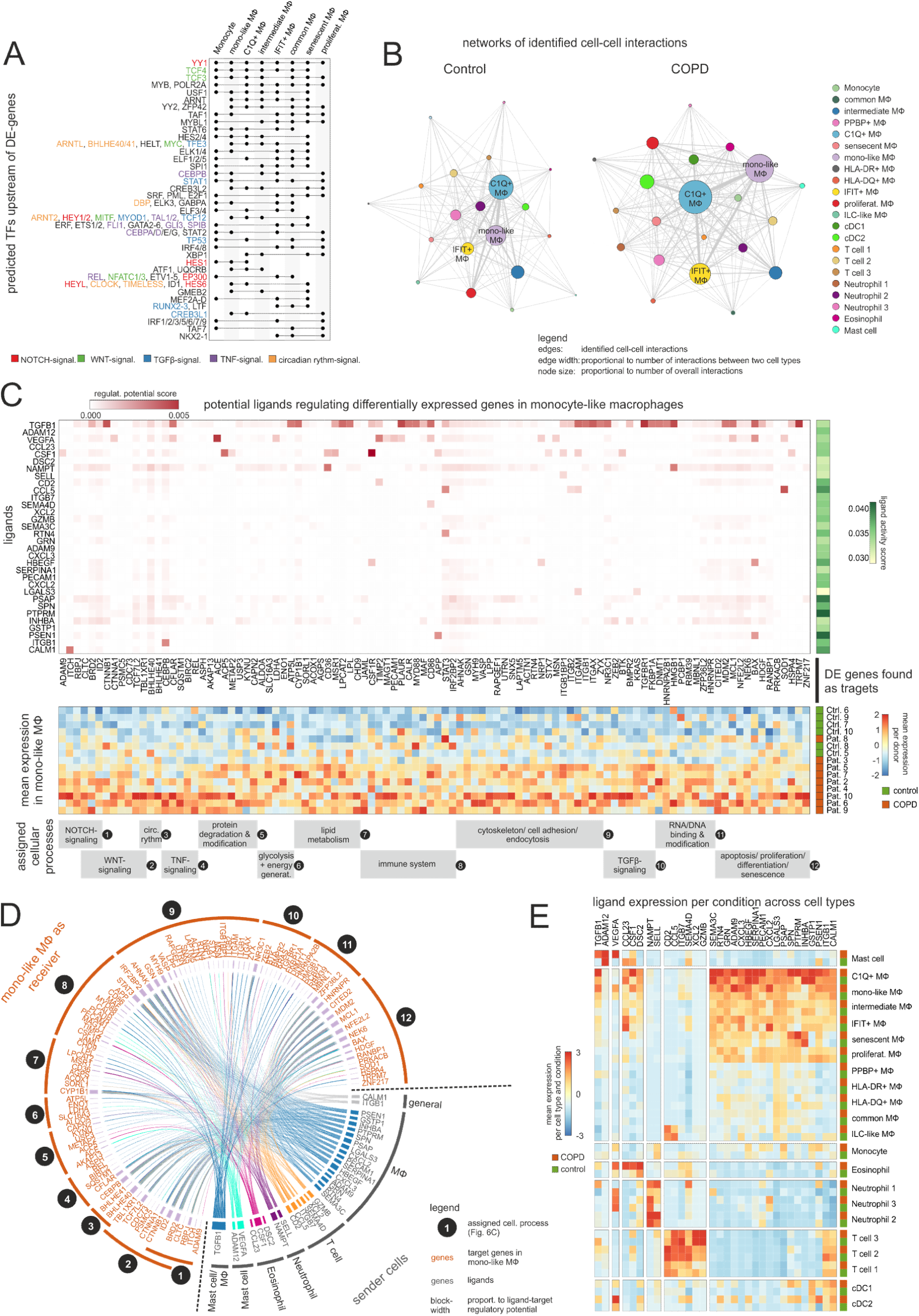
Modelling the cell-to-cell interactions of BALF cells. **(A)** UpSet plot of predicted transcriptional regulators of DE genes. Dots indicate which clusters contain and share predicted transcriptional regulators. The names of selected regulators are shown on the right side of the plot with the font color indicating the association with NOTCH, WNT, TGF-β, TNF or circadian rhythm signaling. **(B)** Network representation of predicted cell-to-cell interactions derived from CellPhoneDB. The names of the three most interconntected cell types are displayed. **(C)** The top heat map represents the NicheNet analysis for mono-like macrophages showing the potential of predicted upstream ligands (on the y-axis) to regulate downstream targets derived from DE genes (on the x-axis). The ligand activity score is depicted as the color-coded area under the precision recall curve (AUPR) on the right side of the plot. The ligands were grouped and ordered based on the cell types from which they are most likely expressed (acc. to Fig. 6D+E). The mean gene expression of the target genes across donors is shown in the lower heat map. The mean gene expression per donor is represented as a z-transformed value (across all donors). Rows of the heat map are sorted by hierarchical clustering. Target genes are ordered and grouped according to cellular functions and signaling pathways, as illustrated at the bottom of the plot. **(D)** Circle plot showing possible regulatory connections between the ligands expressed on different sender cells (acc. to (E)) and the downstream target genes. Target genes are ordered according to (C) as indicated by the numbering. **(E)** Heat map representation of the mean expression of predicted upstream ligands (acc. to (C)) in controls and COPD patients across cell types in BALF. Columns and rows are sorted by hierarchical clustering. AM = alveolar macrophage; signal. = signaling; TF = transcription factor; MΦ = macrophage; regulat. = regulatory; DE = differentially expressed; Pat. = patient; Ctrl. = control; generat. = generation; cell. = cellular; mono = monocyte; DC = dendritic cell; ILC = innate lymphoid cell.

The major pathways regulating the identified DE genes (NOTCH, WNT, TGFβ, TNF) suggested microenvironmental signals to be the major drivers of transcriptional alterations. We therefore applied a recently introduced model for cell-to-cell interactions based on known receptor-ligand interactions (CellPhoneDB (Efremova et al., 2019, 2020; Vento-Tormo et al., 2018)). Network construction of the cell-to-cell interactions within control samples revealed mono-like and C1Q^+^ macrophages to be the major network hubs (**Figure 6B**). In COPD, overall, predicted cell-to-cell communication was increased, which was particularly obvious for three macrophage states, namely C1Q^+^, IFIT^+^ and mono-like macrophages (**Figure 6B, S6A**). Visualization of major receptor-ligand pairs for mono-like macrophages (**Figure S6B**) expressing elevated cell-to-cell interactions with a large variety of other cell types and states in COPD (**Figure S6A**) revealed several receptor-ligand combinations associated with the TNF superfamily (**Figure S6B**). Furthermore, we found an increased likelihood of interaction between the ligand *TGFB1* and the receptor *TGFBR1* in COPD.

While CellPhoneDB predicts potential receptor-ligand interactions based on their expression on sender and receiver cells, it does not model the downstream transcriptional effects of these interactions. To integrate these downstream effects, we applied the NicheNet algorithm (Bonnardel et al., 2019; Browaeys et al., 2019) and focused the analysis on mono-like macrophages. First, we performed ligand activity analysis to prioritize ligands that could best predict the defined DE genes in mono-like macrophages (**Figure 6C and S6C**). Among the top-ranked ligands, we found *TGFB1*, and the set of identified DE genes, which might be regulated by the predicted upstream ligands, could be assigned to cellular processes that we already found in the aforementioned analysis, such as lipid metabolism, immune system process and cell adhesion and differentiation. Noteworthy, some of these DE genes were also associated with cell signaling pathways containing TFs that were predicted as potential upstream transcriptional regulators (**Figure 6A**), such as NOTCH signaling (*RBPJ* and *ADAM9*) and circadian rhythm (*BHLHE40/41*). The highly complex and interwoven network of ligands and their target genes was displayed in a Circos plot (**Figure 6D**) depicting an overview of the potential sender cells of the top-ranked ligands. Thus, mast cells and macrophages were revealed as sender cells of *TGFB1.* Further analysis showed that specific macrophage subtypes, especially mono-like and C1Q^+^ macrophages expressed *TGFB1*, and that in COPD the expression was increased in these cells and mast cells (**Figure 6E**). In addition to elevated TGFB1 expression in COPD, the receptors with the highest predicted interaction potential score for *TGFB1* (*TGFBR1* and *TGFBR2*) exhibited also higher expression in mono-like macrophages from COPD patients (**Figure S6D**). To assess whether the increase in *TGFB1* expression is translated into an elevated protein amount, we examined the BALF of COPD patients and control donors for the latency-associated peptide TGF-β1 (LAP TGF-β1), which served as a surrogate for TGF-β1 protein levels. This analysis showed a clear tendency towards increased LAP TGF-β1 levels in COPD (**Figure S6E**). Finally, we visualized NicheNet-predicted signaling and transcriptional regulation events between TGF-β1 and its putative target genes shown to be DE in COPD (**Figure S6F**). The nodes in the constructed path were colored according to the expression fold change (FC) between COPD and control. Among the transcriptional regulators were the classical TGF-β signaling mediators *SMAD3* and *SMAD4*, with *SMAD4* showing increased expression in COPD (**Figure S6F**). However, we also found some TFs already predicted as potential upstream transcriptional regulators of DE genes in AMs and that showed increased FCs, such as *EP300* and *MYC*, which are also involved in other signaling pathways and which again illustrate the complex interconnected DE-gene regulation.

In summary, within a highly complex network of transcriptional regulation, we predicted TGF-β signaling as a prominent regulator in mono-like macrophages but also other pathways including NOTCH and TNF signaling.

### The AM pool in COPD is supplied by blood monocytes

Some of the identified pathways regulating the DE genes in mono-like macrophages (*e.g.* TGF-β and NOTCH signaling) are known orchestrators of cell differentiation. Tissue macrophage replenishment has been linked to the proliferation of tissue-resident cells (Hashimoto et al., 2013) but also influx and subsequent differentiation of monocyte-derived cells from the circulation (Guilliams and Scott, 2017). To model the potential contribution of both mechanisms in the human alveolar space and to determine whether there are alterations in COPD patients, we first compared the relative differences of the AM states within the alveolar space (**Figure 7A**). Clearly, the AM states characterized as mono-like and proliferating AMs were increased in COPD, while HLA-DR^+^ and HLA-DQ^+^ AMs were reduced, indicating that replenishment of AMs in COPD might be due to both, tissue-intrinsic proliferation as well as cell influx. The mono-like AMs have transcriptional similarities to monocytes (**Figure 4A**) and at the same time exhibit typical signaling pathways of cell differentiation (**Figure 6A-E**) and thus might originate from monocytes. To further corroborate that the mono-like AM state represents an early stage of monocyte-to-macrophage differentiation, we used a gene signature of murine monocyte-derived macrophages (MDM) from the lungs of smoke-exposed mice (Wohnhaas, Baum, *unpublished data*) and assessed the enrichment of orthologous genes in the human AM states (**Figure 7B**). The strongest enrichment of the MDM signature was found in mono-like and C1Q^+^ AMs. In addition, the investigation of orthologous gene signatures derived from murine lipid-associated macrophages (LAM), which were shown to be monocyte-derived by lineage tracing and also detected in human adipose tissue (Jaitin et al., 2019), again revealed the strongest enrichment in mono-like and C1Q^+^ AMs (**Figure 7B**). The same was also true for orthologous genes derived from Trem2^+^ foam cells of atherosclerotic plaques in mice (Kim et al., 2018) (**Figure 7B**), which were found by Lin et al. to originate from monocytes (Lin et al., 2019), supporting the hypothesis that mono-like and C1Q^+^ AMs are actually derived from monocytes.

**Figure 7.**
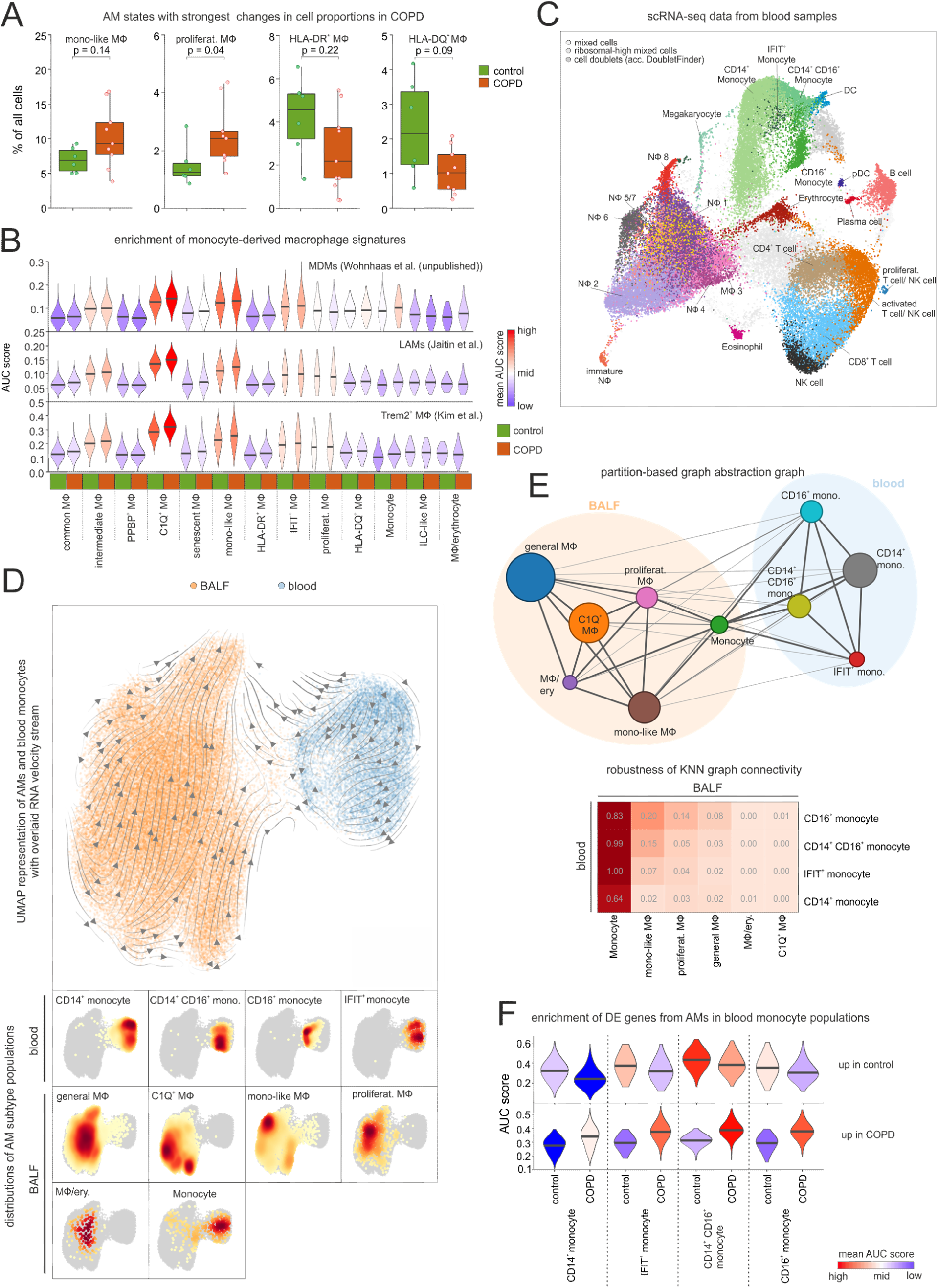
Assessing the relationship between blood monocytes and BALF macrophages. **(A)** Box plots of the relative proportion of macrophage subtypes with representation of individual donors (control n = 6, COPD n = 9). **(B)** Violin plots displaying enrichment of human orthologues of murine monocyte-derived macrophage signature genes across AM states in COPD and control based on ‘Area Under the Curve’ (AUC). **(C)** Integrated scRNA-seq data of blood immune cells annotated according to the four-step annotation approach (acc. to Fig.2A). **(D)** UMAP of embedded macrophages/monocytes from BALF and blood monocytes. Inferred main average vector flow is indicated by velocity streamlines that are projected as vectors. Locations of the main cell types (acc. to the combined labels from Fig. S7A) in the UMAP are indicated by the heat maps at the bottom. **(E)** PAGA graph derived from embedded BALF and blood data (acc. to (D)). The weight of an edge, which reflects a statistical measure of connectivity, is represented as the edge width. The table below summarizes the results of the PAGA connectivity calculation, where a value of 1 indicates a strong connection and 0 indicates a weak connection between two cell types. **(F)** Violin plots displaying enrichment of AM-related DE genes (acc. to Fig.4C+D) in blood monocytes based on AUC. BALF = bronchoalveolar lavage fluid; AM = alveolar macrophage; mono = monocyte; MΦ = macrophage; NΦ = neutrophil; proliferat. = proliferating; MDM = monocyte-derived macrophage; LAM = lipid-associated macrophage.

To establish a direct link from circulating monocytes to the monocyte-related AMs, we performed scRNA-seq of blood immune cells (n = 54,569 cells) (**Figure 7C**) from the same donors from whom the scRNA-seq data of alveolar space immune cells were obtained, and subsequently annotated the cells according to the four-step approach described above (**Figure 2A**). Within the generated dataset, we identified the three known blood monocyte populations comprising classical monocytes (CD14^+^ monocytes), intermediate monocytes (CD14^+^CD16^+^ monocytes) and non-classical monocytes (CD16^+^ monocytes) along with a small monocyte population that expressed high numbers of interferon-associated genes (IFIT^+^ monocytes) (**Figure 7C**). We next described the relationship between blood-derived monocytes and alveolar space-derived monocytes and macrophages by building a model to determine which of the monocyte subtypes in the blood would most likely give rise to the mono-like AM state. For this purpose, we combined the blood and BALF data while considering donor batches by performing a joint embedding based on highly variable genes that are shared across patients (**Figure 7D**). While this approach enabled the combination of the blood and alveolar space data, we observed a reduced resolution of the defined AM states and therefore continued with a simplified annotation for the analysis of the embedded data (**Figure 7D, S7A**). Projection of RNA-velocity vectors calculated by the scVelo method (Bergen et al., 2019) in a batch-corrected manner onto the embedded data (**Figure S7B**) and the inference of the main average vector flow visualized by velocity streamlines (**Figure 7D**) revealed a clear motion of blood monocytes towards the AMs, further supporting circulating monocytes to be precursors of AMs in the alveolar space. Since RNA velocity visualization on the UMAP did not reveal a clear link between individual AM states and blood monocyte subsets, we calculated a higher-order representation using partition-based graph abstraction analysis (PAGA) (Wolf et al., 2019) (**Figure 7E**). The strongest connection was derived between blood monocytes and monocytes identified in the alveolar space. To evaluate the connectivity of the PAGA network more precisely, we used the connectivity matrix as a test statistic to define the highest likelihood for each of the blood monocyte subtypes to be related to the different AM states in the alveolar space (**Figure 7E**). The monocytes within the alveolar space served as positive controls indicating very high relationships. However, importantly, we could establish the strongest connections between the CD16^+^ monocyte subtype in blood and the mono-like AMs in the alveolar space, further supporting that the mono-like AMs are most likely an early functional state of AMs after circulating monocytes enter this tissue compartment.

Lastly, we assessed whether the blood monocytes possessed transcriptional differences between COPD and control. Applying GO-shuffling analysis to immune cells of the blood identified numerous gene sets that separated COPD patients from controls, whereby CD16^+^ monocytes contained the largest number of immune-related terms (**Figure S7C**). Intriguingly, some of these gene sets were associated with cellular extravasation and leukocyte adhesion to vascular endothelial cells, which was also reflected by the heat map representation of the genes upregulated in COPD (*e.g.* including integrins *ITGB1, ITGB2, ITGA4*, and *ITGAL*) (**Figure S7D**). We further investigated whether the DE genes in AMs from COPD patients were already altered in blood monocytes (**Figure 4C-G**). Clearly, these genes were altered in the different blood monocyte subtypes derived from COPD patients with CD14^+^CD16^+^ and CD16^+^ monocyte subtypes showing the strongest enrichment of AM DE genes upregulated in COPD (**Figure 7F**). Of particular interest, MHC class I and II genes were found to be expressed at lower levels in COPD-derived monocytes, while mitochondrial genes were upregulated, supporting a systemic component of COPD leading to transcriptional changes in circulating immune cells such as monocytes (**Figure S7E**).

In summary, we provide evidence that blood-derived monocytes contribute to the AM pool, that this contribution is elevated in COPD patients, and that blood monocytes already show transcriptional changes reminiscent of those observed in cells from the AS strongly arguing for a systemic component in COPD.

## Discussion

There is a global spread of COPD with increasing incidence, prevalence, morbidity and mortality mainly due to increasing air pollution and high global smoking rates (Rabe and Watz, 2017). Yet, the cellular and molecular mechanisms of this heterogeneous disease syndrome are far from being fully understood (Rabe and Watz, 2017). Not surprisingly, the diagnosis of COPD is solely based on clinical parameters due to the lack of molecularly defined biomarkers and, as a consequence, causal therapies are lacking because of incomplete understanding of the complex pathophysiology (Barnes et al., 2015). Here, we set out to contribute to the understanding of COPD by applying high-resolution single-cell technologies together with sophisticated computational approaches to freshly available biospecimens from COPD patients. We focused on the alveolar space, which was sampled by BALF, and peripheral blood as these are two compartments that are clinically accessible. Furthermore, in this first study, we focused on patients diagnosed with earlier clinical grade disease (GOLD 2) since it has been hypothesized that reprogramming of involved cells as a potential therapeutic option might only be possible at earlier grades (Sun and Zhou, 2019). Single-cell transcriptomes from both alveolar space and blood were reliably obtained using the array-based scRNA-seq technology Seq-Well (Gierahn et al., 2017), while a widely used droplet-based method performed inferiorly on granulocytes, highlighting the need for further adaptation of single-cell technologies to difficult clinical specimens. Since cell-type classification is of particular importance when applying scRNA-seq technologies to patient cohorts, we developed a four-step cell-type annotation approach. This procedure identified all major immune cell types but also subsets, particularly in the AM compartment that is categorized by unexpectedly high transcriptional plasticity. At this point, it is important to note that the term ‘alveolar macrophage’ is a generic term, because although bronchoalveolar lavage mainly involves the collection of cells from the alveolar space, it also involves the collection of some cells from the bronchioles. For simplicity, the macrophage compartment from BALF in this study is referred to as AMs. *In silico* prediction of metabolic changes based on scRNA-seq data suggested alterations in lipid metabolism of the AM compartment, which we validated by lipidomics with significant changes in cholesteryl ester metabolism in COPD. In addition, we also observed alterations in quantities of individual AM states, for example, proliferating and mono-like macrophages were elevated in COPD. Using a newly developed computational approach to define DE genes on single-cell level between AM states in COPD and control individuals, we observed changes in COPD that were shared by most AM states, for example, reduced expression of MHC class I molecules. Transcription factor binding prediction and receptor-ligand interaction modelling and downstream transcriptional signature prediction suggested that especially TGF-β signaling but also other pathways including NOTCH, WNT, and TNF-signaling were elevated in COPD. PAGA and RNA velocity analysis of cells from the peripheral blood and the alveolar space, in a patient cohort setting, suggested that a proportion of the AM pool is replenished from the systemic monocyte pool circulating in peripheral blood. Furthermore, COPD-related signatures derived from cells in the alveolar space were already enriched in the peripheral blood monocyte pool, particularly in CD14^-^CD16^+^ and CD14^+^CD16^+^ subsets, clearly indicating that the pathophysiology of COPD is not restricted to the lung.

The COPD study presented here required adaptation, optimization and new development of experimental and computational procedures to derive clinically meaningful results from precious and fragile biospecimens. When comparing with MCFC, the array-based scRNA-seq method Seq-Well outperformed one of the droplet-based methods in capturing fragile granulocytes, a finding that was recently also observed by others (Travaglini et al., 2019). Another important factor in the investigation of clinically relevant biospecimens using scRNA-seq is the correct annotation of cell types, which currently presents a major problem in the field (Abdelaal et al., 2019). Recently, machine learning-based methods were successfully used for cell type annotation of single-cell data (Alquicira-Hernandez et al., 2019; Hou et al., 2019; Lopez et al., 2018; Song et al., 2019) and – surprisingly - it was reported that the machine-learning methods themselves hardly represented a qualitative difference, as most supervised methods performed similarly well (Abdelaal et al., 2019; Köhler et al., 2019). Still, a main issue seems to be the access to well-curated training data. To address these limitations, we combined several prior knowledge methods based on marker genes or reference expression profiles to obtain multiple labels per cell. These multi-labeled cells were then used in a cross-validation approach to train a gradient boosting classifier and predict the cell type of individual cells and thereby consolidate the former labels. In addition to the expression values, we included additional features and iteratively curated the initial labels. Using this approach, all cells were individually annotated in a reproducible and comparable manner, although no definitive ground truth was initially available.

Another still existing challenge in SCG is the identification of DE genes within a cohort setting, here COPD and controls. Current methods for DE gene calculation are regularly performed after generating ‘mini-bulk’ gene expression values for defined clusters of cells, which are then used for comparison of two groups, *e.g.* COPD and controls. While computationally straightforward, this approach dismisses the single-cell gene information. Furthermore, appropriate significance levels are difficult to determine for these tests as there are various sources of variability between biological samples, which are difficult to emulate *in silico*. To address these limitations, we developed a novel DE analysis approach that exploits the single-cell measurements but avoids distribution assumptions and determines the significance of the observed effects using a permutation test. Such tests require larger parameter spaces and therefore have not been considered in benchmark studies for multi-sample comparisons (Crowell et al., 2019). However, the sufficiently large number of individuals included in our study facilitated the application of such a method (Ernst, 2004).

By applying these computational innovations, we discovered new insights into the pathophysiology of COPD. We identified altered lipid metabolism in AMs of COPD GOLD 2 patients, which was characterized by increased accumulation of cholesteryl ester. All tissue macrophages exhibit ‘accessory’ functions to support parenchymal cells (Okabe and Medzhitov, 2016), with AMs regulating the surfactant homeostasis in the lung (Remmerie and Scott, 2018). Thus, they are highly associated with lipid metabolism. Failure of this ‘accessory’ function is reflected by pulmonary alveolar proteinosis (PAP), in which defective GM-CSF signaling in AMs causes an accumulation of surfactant-derived lipids in AMs and the AS (De Aguiar Vallim et al., 2017; Sallese et al., 2017; Trapnell et al., 2019). However, due to the expression of *ABCG1* in AMs from COPD patients, the accumulation of cholesteryl ester must be due to alternative molecular mechanisms than in PAP (De Aguiar Vallim et al., 2017; Baker et al., 2010). There might be a link to cholesterol accumulation in macrophages in atherosclerosis, as statin treatment of patients with atherosclerosis and COPD was shown to result in reduced exacerbations and mortality of COPD in some of these patients (Ingebrigtsen et al., 2015).

Other aspects of pathophysiology such as monocyte recruitment might also be shared between the two diseases. A recently proposed model (Guilliams and Scott, 2017) suggested that under homeostatic conditions survival of tissue-resident macrophages is supported by self-renewal within the local microenvironment while monocyte recruitment is rather limited (Scott et al., 2016). During inflammation, tissue-resident macrophages retain the ability to self-renew, but at the same time blood-derived monocytes are recruited (Hashimoto et al., 2013). In COPD, we found evidence for both local proliferation of a small fraction of AMs and recruitment of blood monocytes. Indeed, mono-like and C1Q^+^ AMs exhibited strong enrichment of monocyte-derived macrophage signatures and RNA velocity analysis supported a differentiation process between blood monocytes, particularly the CD16^+^ subset and the mono-like cell state within the AM space. These findings are further corroborated by previous findings demonstrating that murine CD16^+^ blood monocytes can differentiate into lung macrophages (Schyns et al., 2019). Another line of evidence comes from murine smoking models illustrating that Cx3cr1^+^ macrophages expand in response to cigarette smoke (Lee et al., 2012) and exhibit a profibrotic function (Aran et al., 2019a). It will be interesting to determine whether Cx3cr1^+^ cells in atherosclerosis associated with Ly6c^-^ monocytes, the murine counterpart of human CD16^+^ monocytes (Moore et al., 2013), are similar to the Cx3cr1^+^ macrophages in the lung.

Our pathway prediction identified elevated NOTCH, WNT, TGF-β, and TNF signaling, which also supports an increased differentiation of monocytes into AMs as this process has been recently associated with TGF-β (Yu et al., 2017) and NOTCH signaling (Bonnardel et al., 2019). Elevated NOTCH signaling in alveolar epithelial cells has been implicated in chronic respiratory diseases, such as pulmonary fibrosis (Reyfman et al., 2019) while WNT signaling seems to be downregulated in human epithelial cells from COPD patients, leading to a decrease in their repair capacity (Skronska-Wasek et al., 2017). In macrophages, elevated levels of NOTCH ligands such as DNER have been associated with respiratory diseases, and NOTCH signaling was shown to be upstream of IFN-γ induction in cigarette-exposed animals (Ballester-López et al., 2019; Reyfman et al., 2019) as well as in COPD patients (Ballester-López et al., 2019). Of interest, SNPs in DNER have been associated with a higher risk of COPD (Busch et al., 2017; Hancock et al., 2012). While crosstalk between NOTCH, WNT, and TGF-β pathways are known for non-small cell lung cancer (Li et al., 2011; Ohnuki et al., 2014), this has not been reported yet for COPD. Increased expression of TGFB1 was reported for COPD (de Boer et al., 1998) and single-cell level analysis clearly illustrates that in the alveolar space only some immune cell states, here mast cells, mono-like and C1Q^+^ macrophages are contributing to the altered expression levels.

The predicted pathways might also be involved in immunosenescence in COPD. NOTCH, WNT and TNF signaling can induce mTOR signaling (Chan et al., 2007; Saxton and Sabatini, 2017), which was recently associated with cellular senescence in lung cells (Houssaini et al., 2018). As COPD develops preferentially in elderly people (Halbert et al., 2006) often with several comorbidities, cellular aging might be a hallmark of the disease (Barnes, 2017). Features of cellular senescence comprise an increase in the number of mitochondria and mitochondrial dysfunction, which is reflected by increased proton leakage and an associated increase in ROS production (Gorgoulis et al., 2019; Korolchuk et al., 2017). Oxidative stress due to increased ROS production is a well-known feature of COPD and there is evidence that this is partly due to mitochondrial dysfunction (Bowler et al., 2004; L et al., 2012; McGuinness and Sapey, 2017; Ryter et al., 2018). In line with cellular senescence, we found increased expression of mitochondrial genes and increased proton leakage in the mitochondria of AMs from COPD patients.

Additionally, the reduced chemotactic capacity of AMs in COPD might also be a result of aged immune cells (Oishi and Manabe, 2016; Shaw et al., 2013; Solana et al., 2012). Dysregulation of macrophage chemotaxis has also been described as a consequence of smoking (Berg et al., 2016) and in lung cancer (Lemarie et al., 1984). Reduced migratory capacity of AMs can have deleterious consequences for the lung, as it reduces the efficient removal of pollutants from the alveolar space, which can lead to cell death and the induction of inflammation. Moreover, the clearance of the alveolar space is further deteriorated due to decreased phagocytosis in AMs in COPD (Taylor et al., 2010).

Further dysregulation of macrophage function in COPD was documented by significant downregulation of molecules associated with antigen presentation, especially MHC class I. This finding is in accordance with previous studies linking down-regulation of surface MHC class I in COPD with impaired immunoproteasome activity (Hodge et al., 2011; Kammerl et al., 2016). Downregulation of MHC class I can trigger NK-dependent killing, but the simultaneous expression of the non-classical MHC class I molecule HLA-E on AMs likely protects immunosenescent AMs from effective clearance by NK cells (Pereira et al., 2019). AMs expressing MHC class I at low levels are less efficient in inducing an antiviral immune response, which may explain the high susceptibility of COPD patients to viral infections, one of the main reasons of disease exacerbations (Woodhead et al., 2005). Reduced MHC expression was also observed on circulating blood monocytes, which further underlines the systemic character of COPD (Agusti, 2005; Agusti and Soriano, 2008; Fabbri and Rabe, 2007). Collectively, we provide a starting point for a systems-level understanding of COPD pathophysiology based on information derived from individual cells in two major tissue compartments, the alveolar space and peripheral blood. The next steps will encompass data integration derived from cells within the lung parenchyma, spatial information about different areas within the lung, scaling towards studies including additional patients also at additional disease grades, and if possible in a longitudinal setting to determine the overall patient-to-patient heterogeneity of the disease. To be able to continuously integrate future single cell-level information, we provide data and analyses in an integrated fashion on https://www.fastgenomics.org/, as this web portal allows seamless integration of additional datasets.

While we recognize that an explorative study as the one reported here cannot yet provide a definitive list of potential biomarkers to be further tested in larger clinical trials, our findings already indicate some principles. First, we did not identify single markers that are specific for COPD GOLD 2, suggesting that signatures rather than individual markers will have to be developed. For example, the loss of MHC class I molecules on myeloid cells is not only seen in COPD, it is also described in cancer (Noy and Pollard, 2014) and in influenza (Koutsakos et al., 2019). Similarly, the increase of the mono-like macrophage cell state as a hallmark for cell influx from the systemic circulation will also be seen in other pathophysiological situations of the lung, for example, acute infections such as pneumonia (Goto et al., 2004) or influenza infection (Aegerter et al., 2020). In contrast, the combination of these changes together with the changes observed in lipid metabolism or the COPD AM signature also observed in peripheral blood monocytes are good starting points for future biomarker development for COPD diagnostics and outcome prediction.

In summary, we provide clear evidence that the application of improved computational approaches combined with a clinically applicable scRNA-seq technology to two major tissue compartments in COPD and control patients leads to a deeper understanding of the complex pathophysiology and the involvement of the immune system in this devastating disease, which is a prerequisite for the development of new diagnostic biomarkers as well as causal therapies. Providing the data and our computation of the data in an integrated fashion on a publicly available web portal is a starting point to include future data generated from additional patient samples and tissue compartments involved in human COPD.

## Acknowledgments

We thank F. Gondorf for technical assistance with Seahorse, T. Quast and K. Zölzer for their support in using microscopes, and S. Mukherjee and B. Taschler for their support in the development of GO-shuffling.

## Author contributions

Conceptualization, K.B. D.S., P.B., and J.L.S; Methodology, K.B., S.W.H., J.L.S., B.R., F.B., H.D., J.H., M.L., and F.J.T.; Software: K.B., S.W.H., B.R., E.D., and M.L.; Investigation, K.B., W.F., T.S.K., A.H., B.R., E.D., M.L., N.R:, C.O.S., S.W.H., L.B., P.G., M.B., K.H., H.T., M.K., H.J.G.F., J.S.S., E.H., C.T., A.F., D.T., A.C.A., and T.U.; Biospecimen/ enzyme resources, C.P., T.S., D.S., I.H.K. and M.G.; Writing – Original Draft, K.B. and J.L.S.; Writing – Review & Editing, K.B., J.L.S., W.F., T.S.K., A.H., B.R., E.D., M.L., N.R., C.O.S., S.W.H., A.W., L.B., P.G., M.B., C.T.H., M.H.W., T.K.H., J.S.S., E.H., I.H.K., M.G., C.T., A.K.S., H.D., M.B., P.B., N.Y., A.C.A., T.U., J.H., F.J.T., D.S.; Visualization, K.B., A.H., N.R., A.C.A., and M.L.; Supervision, J.L.S.; Project Administration, J.L.S.; Funding Acquisition, W.F., J.L.S., F.J.T., J.H., N.Y., A.K.S., M.G., and M.B.

## Data and materials availability

Data will be deposited at the European Genome-phenome Archive (EGA) and will be accessible via the EGA study-ID ‘EGAS00001004369’. In addition to data deposition on EGA, we provide an interactive platform for data inspection and analysis via FASTGenomics. The FASTGenomics platform provides also a normalized count table of the Seq-Well datasets used in this study. We have also deposited the code for the novel DE-analysis approach used in this study on Zenodo (https://doi.org/10.5281/zenodo.3717776).

---

**Figure S1.**
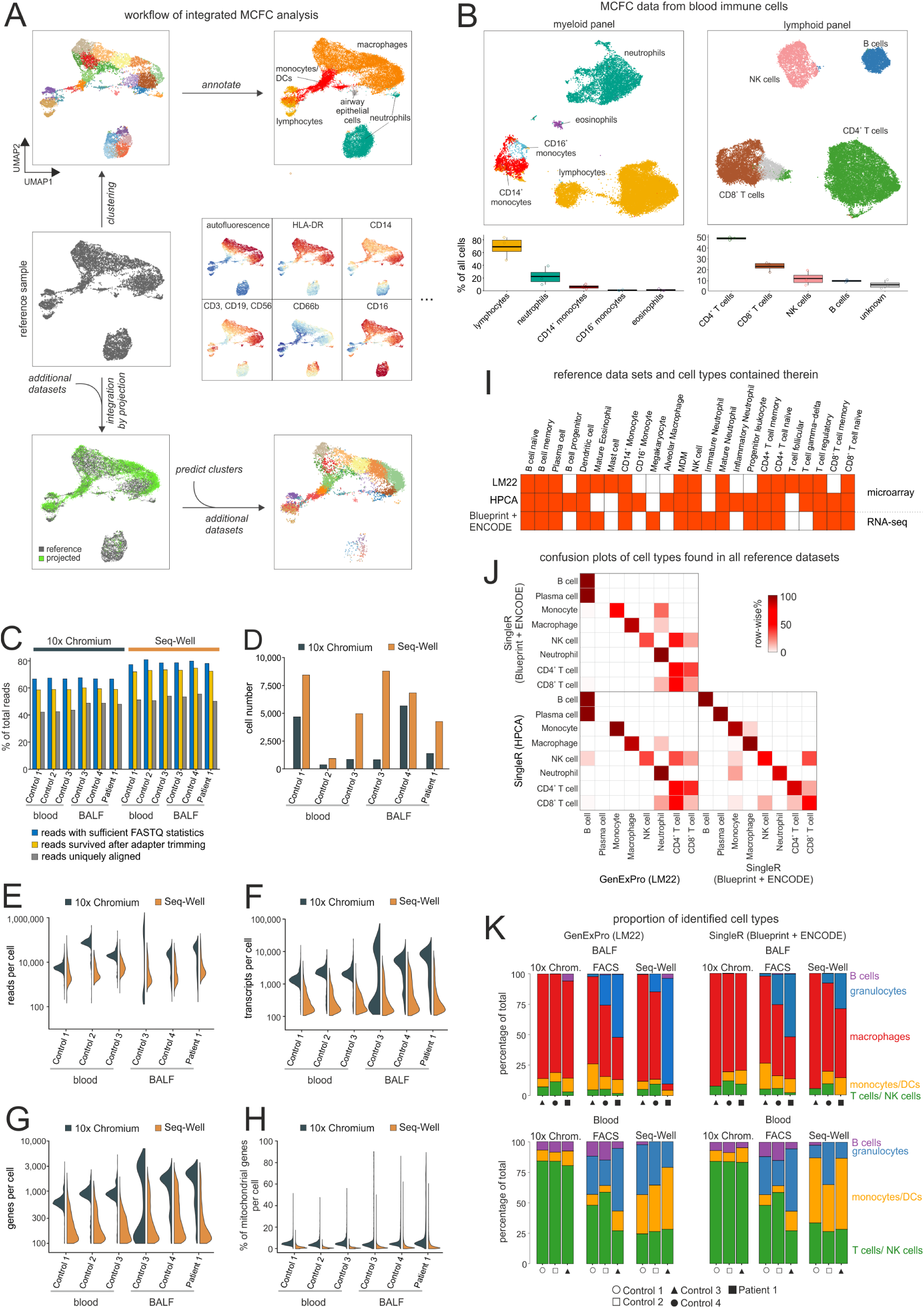
Comparison of scRNA-seq data from Seq-Well and 10x Chromium technology (related to Figure 1). **(A)** Workflow of MCFC analysis obtained from different patients, starting from the UMAP representation of a compensated reference sample, through clustering of the data and annotation of the clusters based on the protein fluorescence intensities. The other samples are then projected onto the UMAP reference sample and the accuracy of the projection is evaluated by predicting and comparing clusters. **(B)** UMAP representation of MCFC data obtained from blood immune cells of three donors. Relative proportion of identified cell types per donor displayed as boxplots. **(C) – (H)** scRNA-seq library statistics of data obtained from Seq-Well and 10x Chromium technology. **(I)** Overview of the cell types contained in the reference files used for cell-type annotation. The orange color indicates that the respective cell type is included in the reference file. **(J)** Confusion plots showing the concordance between the respective cell-type annotations across the different annotation methods. Only cell types that can be found in all reference files as shown in (I) are displayed. **(K)** Stacked bar plots of the relative cell-type proportions for MCFC, which served as ground truth, and cell type-proportions of the two scRNA-seq technologies, as predicted by SingleR with the Blueprint + ENCODE dataset as reference and the novel cell-type annotation method GenExPro. BALF = bronchoalveolar lavage fluid; MCFC = multi-color flow cytometry.

**Figure S2.**
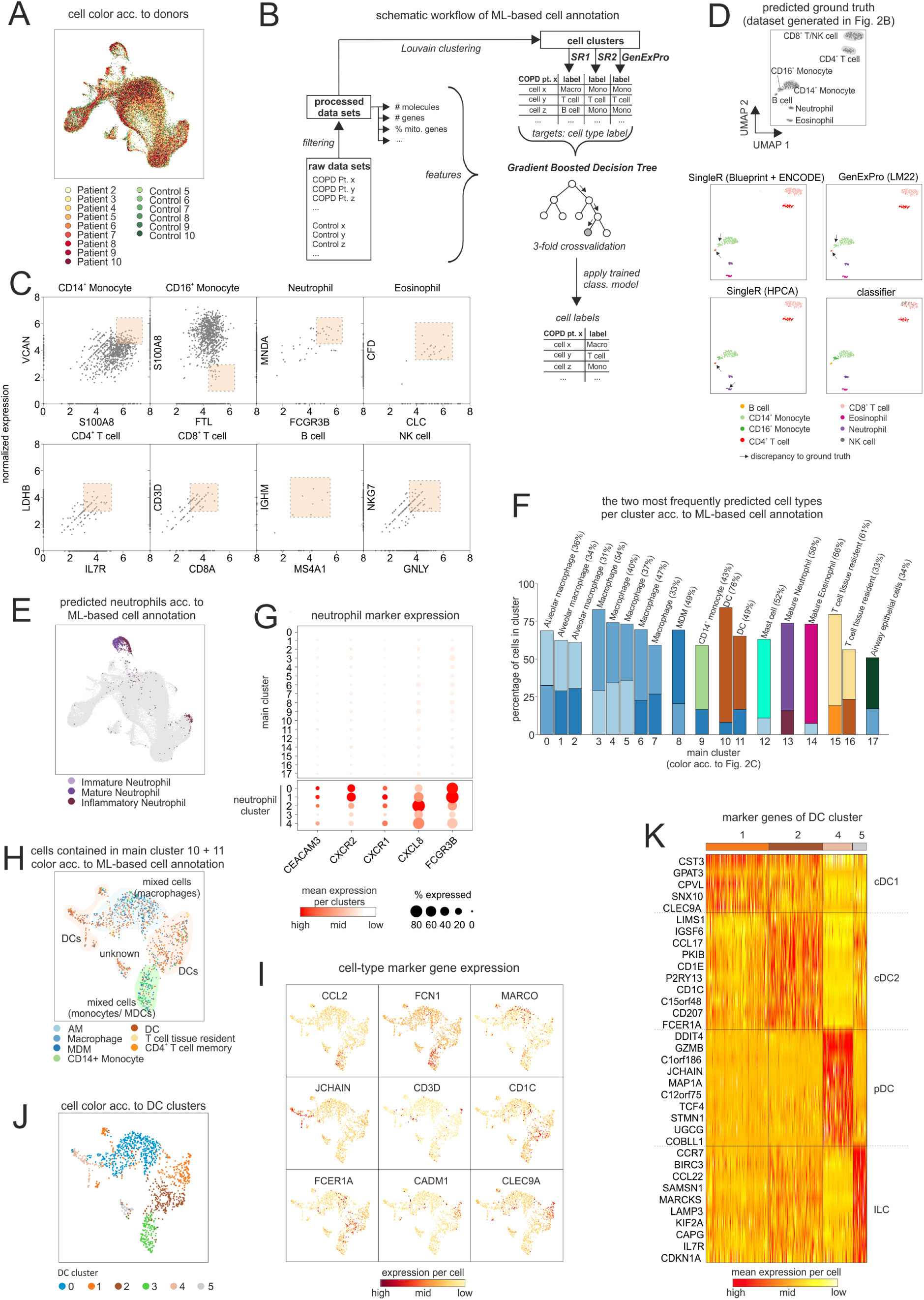
Benchmarking of the four-step approach of BALF celltype annotation (related to Figure 2). **(A)** UMAP representation of the integrated dataset with cell coloring according to the donors. **(B)** Scheme of a gradient boosted decision tree-based machine learning-approach for cell-type annotation **(C)** Generation of a ‘clean’ Seq-Well dataset for benchmarking the machine learning-based cell-type annotation by selecting cells from a blood dataset (patient 6 acc. to Table S1) that express unique markers of a certain cell type. The selected cells, which are combined in a benchmarking dataset, are marked by the dashed rectangles with a reddish background. **(D)** UMAP representation of the benchmarking dataset (acc. to (B)) and coloring of the cells according to the cell annotation methods. The ground truth is derived based on the unique cell-type marker gene expression in (B). Accumulation of cells that are annotated by the respective annotation methods, but show a deviation in the annotation with respect to the ground truth, are marked with an arrow. **(E)** UMAP representation of the integrated dataset with the coloring of different neutrophil states as predicted by the machine learning-based cell-type annotation. Non-neutrophils are colored light gray. **(F)** Bar plot displaying the two most common cell types for each main cluster (acc. to Fig.2B) according to machine learning-based cell annotation, in color-coded form (acc. to Fig.2C). In addition, the most frequently occurring cell type is written above the respective cluster together with its relative frequency. **(G)** Expression of known neutrophil-associated genes represented in a dot plot. Main clusters are shown, with the main cluster 13 subdivided into the neutrophil clusters 0-4 (acc. to Fig.2F). **(H)** UMAP representation of cells contained in main clusters 10 and 11 (acc. to (B)). Coloring according to the machine learning-based cell-type annotation. Accumulation of cells with the same identity were highlighted with colored background and labeled accordingly. **(I)** Feature plots showing the expression of cell type-specific markers (*CCL2* and *FCN1* = monocytes; *MARCO* = macrophages; *CD3D* = T cells; *FCER1A, JCHAIN, CD1C, CADM1* and *CLEC9A* = DCs and DC subtypes)**. (J)** UMAP representation and clustering of the cells contained in main clusters 10 and 11. **(I)** Heat map of markers genes for DC clusters 1, 2, 4, and 5 (acc. to (J)), which were predicted to contain mainly DCs according to the machine learning-based cell-type annotation (acc. to. (H)). Rows of the heat map are sorted by hierarchical clustering. BALF = bronchoalveolar lavage fluid; DC = dendritic cell; AM = alveolar macrophage; Pt. = patient; SR1 = SingleR (HPCA); SR2 = SingleR (Blueprint + ENCODE); Macro = macrophage; Mono = monocyte.

**Figure S3.**
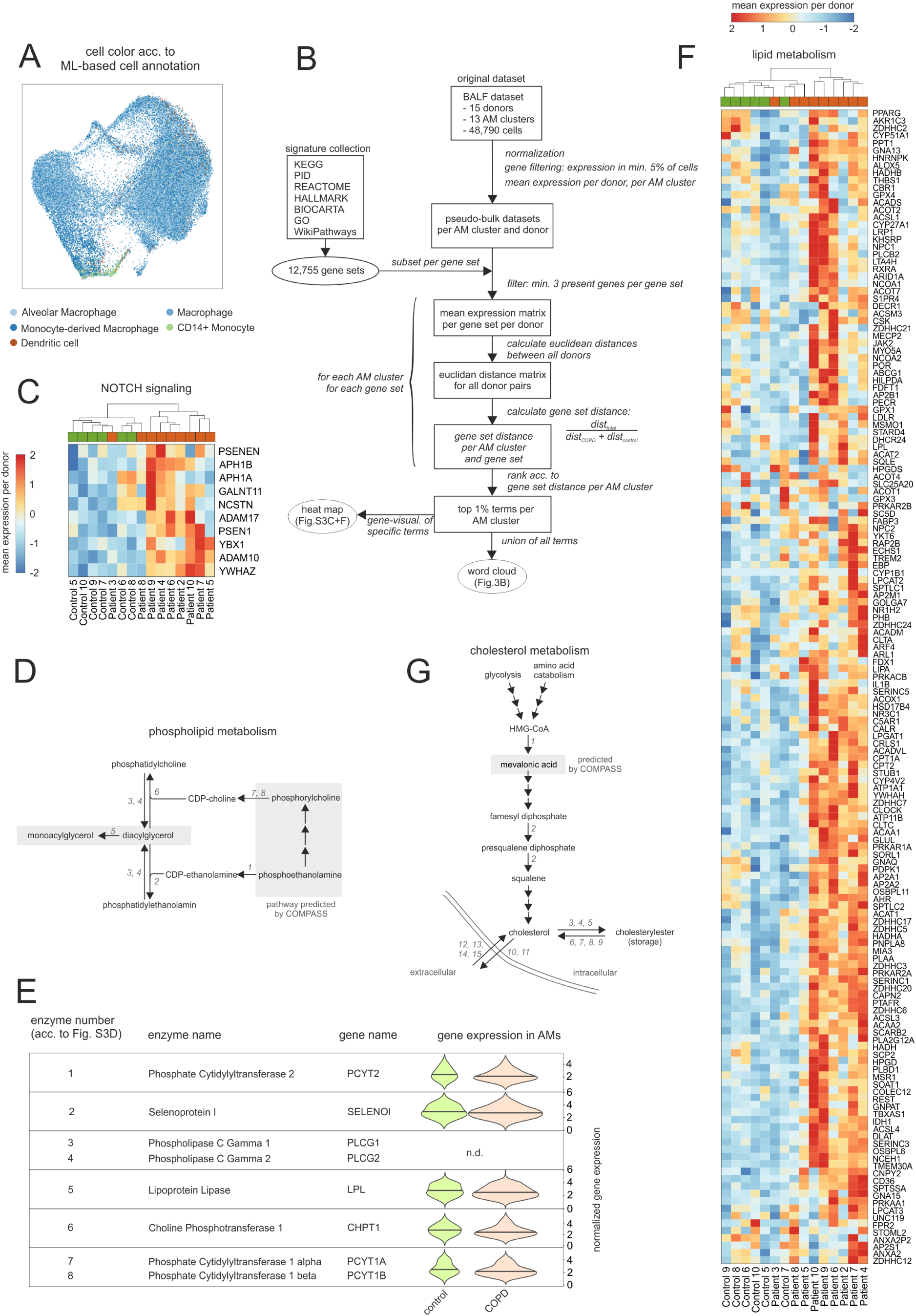
Characterization of altered lipid metabolism in AMs of COPD patients (related to Figure 3). **(A)** UMAP representation and clustering of cells contained in main clusters 0-9, which are annotated as monocytes or macrophages (acc. to Fig. 2B+C). Coloring according to the machine learning-based cell-type annotation. **(B)** Schematic workflow of the GO-shuffling approach. **(C)** Heat map of NOTCH-signaling associated genes predicted by the GO-shuffling approach. The mean gene expression per donor is represented as a z-transformed value (across all donors). Columns and rows of the heat map are sorted by hierarchical clustering. **(D)** Scheme of the part of the phospholipid metabolism containing signaling pathways and metabolites for which Compass predicted a difference between COPD and control. Pathways and metabolites predicted by Compass are highlighted with a gray background. Enzymes involved in metabolism are abbreviated with a number that identifies them also in Fig. 3D. **(E)** Violin plots displaying the gene expression of enzymes involved in phospholipid metabolism (acc. to (D)). The plots show the expression across the donors, whereby the donors were downsampled to the same number of cells, followed by downsampling to the same number of cells between COPD and control. The plots display cells with an expression > 0. The areas of the violin plot are scaled proportionally to the number of observations. **(F)** Heat map of lipid metabolism-associated genes predicted by the GO-shuffling approach. The mean gene expression per donor is represented as a z-transformed value (across all donors). Columns and rows of the heat map are sorted by hierarchical clustering. **(G)** Schema of the key steps in cholesterol metabolism and storage. Metabolites predicted by Compass are highlighted with a gray background. Enzymes involved in metabolism are abbreviated with a number that identifies them also in Fig. 3D. BALF = bronchoalveolar lavage fluid; ML = machine learning; acc. = according; AM = alveolar macrophage; visual. = visualization.

**Figure S4.**
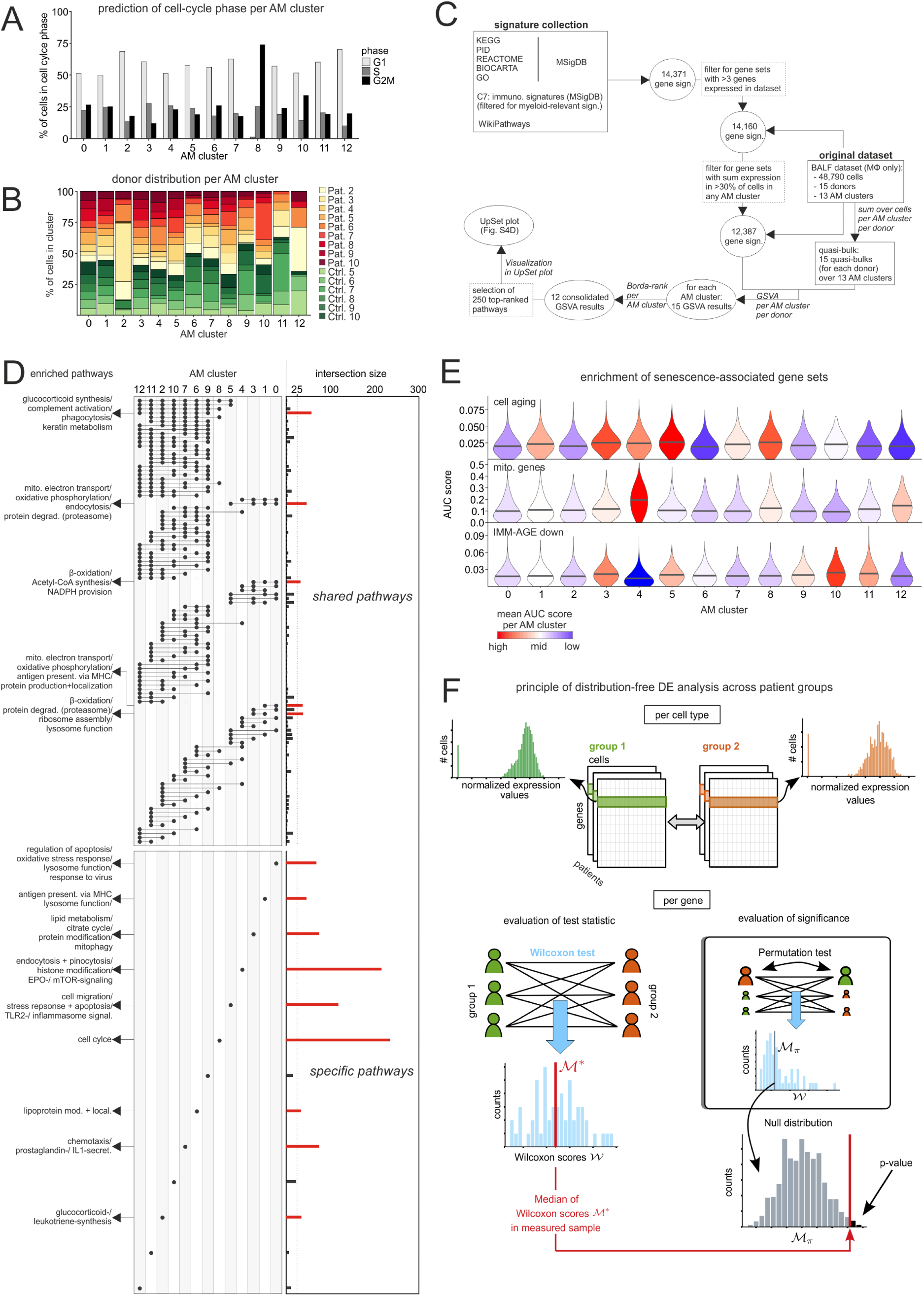
Characterization of identified AM subtypes (related to Figure 4). **(A)** Bar plot representation of the proportion of cells in the respective cell cycle states per cluster (acc. to Fig. 3A). **(B)** Stacked bar plot showing the proportion of individual donors in each cluster. **(C)** Schematic workflow to predict the cellular functions of each cluster based on GSVA. **(D)** UpSet plot of the GSVA results (acc. to (C)). Terms of cellular functions found in the same clusters are grouped into bins and the size of the bins is represented as a bar plot on the right, with bins containing more than 25 terms (dashed line) colored red. On the left side, dots indicate which clusters contain and share the binned terms. Frequently occurring terms of cellular functions within the bins containing more than 25 terms are shown. **(E)** Violin plots displaying enrichment of different gene sets across clusters based on ‘Area Under the Curve’ (AUC). **(F)** Schematic workflow of the permutation test-based DE analysis approach. BALF = bronchoalveolar lavage fluid; Pat. = patient; Ctrl. = control; sign. = signature; MΦ = macrophage; GSVA = gene set variation analysis; mito. = mitochondrial; degrad. = degradation; mod. = modification; present. = presentation; DE = differential expression.

**Figure S5.**
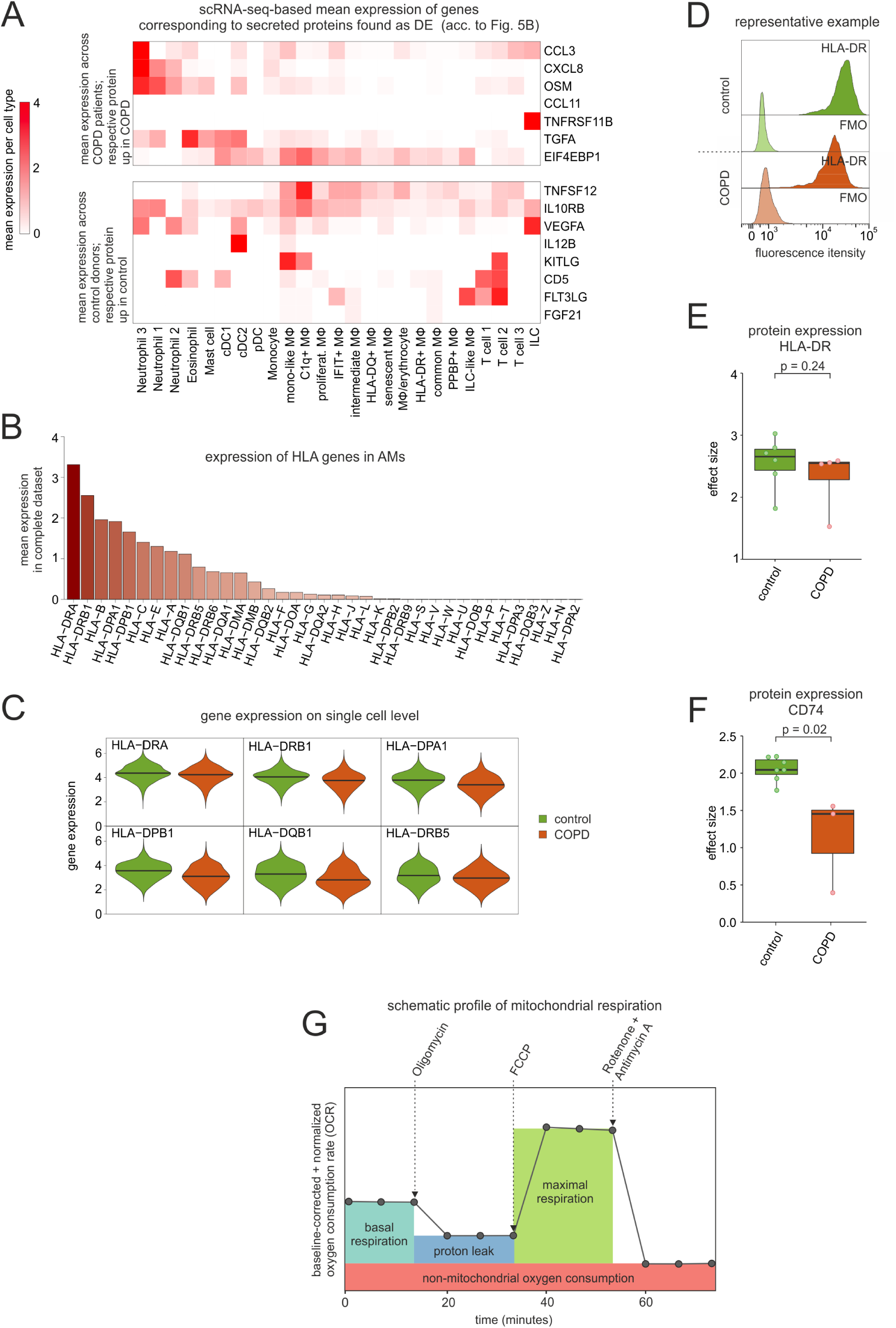
Validation of *in situ* predicted alterations in AMs of COPD patients (related to Figure 5). **(A)** Heat map representing the BALF cell type-dependent expression of genes whose protein counterparts were found in BALF (acc. to Fig. 5B). If the corresponding protein is upregulated in the control group, the mean gene expression was calculated across the control donors and represented as a z-transformed value (across all control donors) (top panel). Similarly, mean expression was calculated only in COPD patients when the corresponding protein is upregulated in COPD (bottom panel). **(B)** Bar plots showing the mean expression of various HLA genes in AMs. **(C)** Violin plots of HLA gene expressions in AMs based on scRNA-seq data. The plots show the expression across the donors, whereby the donors were downsampled to the same number of cells, followed by downsampling to the same number of cells between COPD and control. The plots display cells with an expression > 0. **(D)** Fluorescence intensity histograms showing representative samples of flow cytometric analysis of HLA-DR expression on the cell surface of isolated AMs (FMO = fluorescence minus one). **(E)+(F** Box plots of the calculated effect sizes of HLA-DR and CD74 expression in COPD and control with representation of individual donors (HLA-DR: control n = 6, COPD n = 4; CD74: control n = 7, COPD n = 3). **(G)** Schema of the time-dependent course of the oxygen consumption rate (OCR) and the inferred mitochondrial parameters based on the injection of different compounds (shown at the top of the plot). BALF = bronchoalveolar lavage fluid; acc. = according; MΦ = macrophage; ILC = innate lymphoid cell; DC = dendritic cells; AM = alveolar macrophage; DE = differentially expressed.

**Figure S6.**
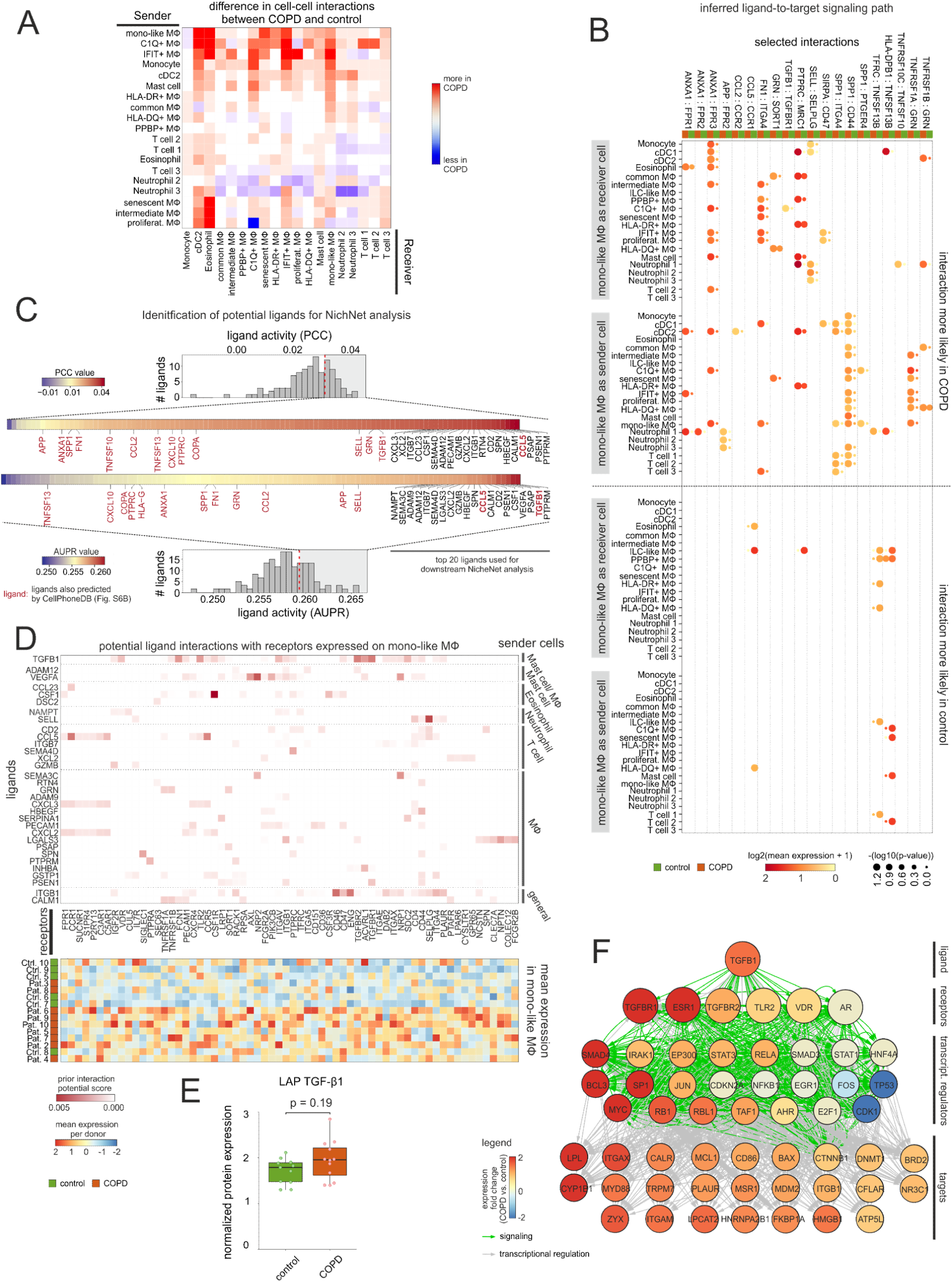
Investigation of cell-to-cell interactions to infer important signaling pathways in AMs (related to Figure 6). **(A)** Heat map representing differences in the cell-to-cell connections between the networks in Figure 6B. Columns and rows of the heat map are sorted by hierarchical clustering. **(B)** Dot plot representation of mono-like macrophage-dependent ligand-receptor interactions predicted by CellPhoneDB that show significant enrichment (represented by the p-value) of the interacting pair in the interacting cell types either in COPD or in the control. Depicted are only selected interactions. **(C)** Illustration of the selection of potential upstream ligands based on the NicheNet analysis. The histograms show distributions based on ligand activity derived from the area under the precision recall curve (AUPR, upper histogram) and the Pearson correlation coefficient (PCC, lower histogram). The ligand activity of the highest-ranked ligands (represented by the grey area in the histograms) is displayed in a color code together with the names of the 20 highest ranked ligands and the ligands also predicted by the CellPhoneDB analysis (highlighted in red). **(D)** The top heat map represents the NicheNet analysis for mono-like macrophages showing the potential of ligands (on the y-axis) interacting with receptors (on the x-axis). The ligands were grouped and ordered based on the cell types from which they are most likely to be expressed (acc. to Fig. 6D+E). The mean gene expression of the receptors on mono-like macrophages across donors is shown in the lower heat map. The mean gene expression per donor is represented as a z-transformed value (across all donors). Rows of the heat map are sorted by hierarchical clustering. **(E)** Box plot of the measured protein expression (by Olink Proteomics) in BALF of LAP TGF-β1 in COPD and control with representation of individual donors (control n = 11, COPD n = 12). **(F)** Representation of inferred ligand-to-target signaling path for TGF-β1 derived from the NicheNet analysis (receptors: acc. to (D); targets: acc. to Fig. 6C+D). AM = alveolar macrophage; MΦ = macrophage; DC = dendritic cell; ILC = innate lymphoid cell.

**Figure S7.**
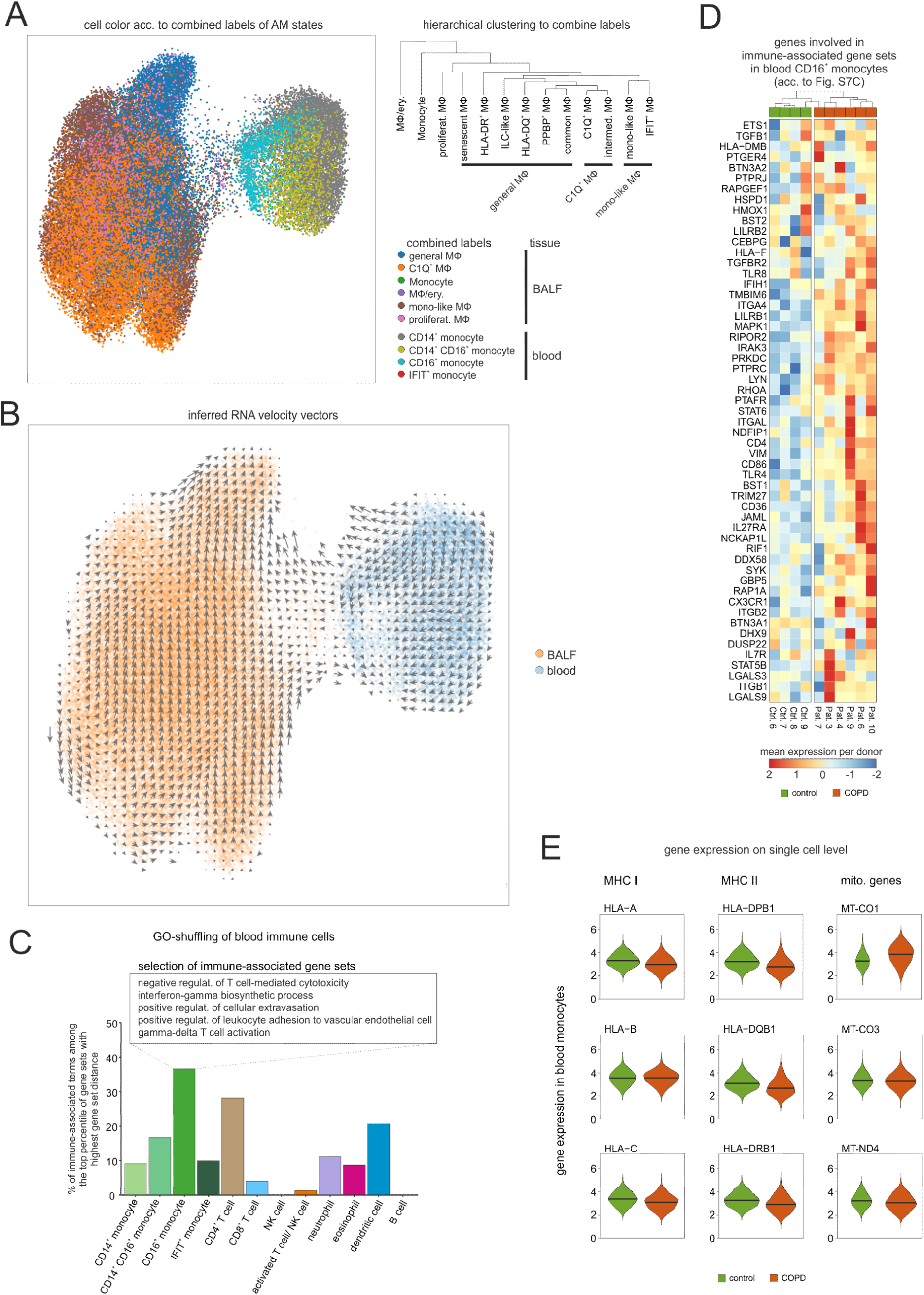
Assessment of immune-related alterations in blood monocytes from COPD patients (related to Figure 7). **(A)** UMAP of embedded macrophages/monocytes from BALF and blood monocytes with coloring according to the cell types derived from the combined labels. The dendrogram on the right side illustrates the transcriptional relationship between the macrophage subtypes and shows how several subtypes were summarized in the combined labels. **(B)** Projection of computed RNA velocity vectors onto the UMAP of the embedded data. **(C)** Bar plot showing for blood immune cell types the proportion of predicted immune-related GO terms within the upper percentile of GO terms with the highest ability to separate COPD patients from control donors according to GO-shuffling. A selection of immune-related GO terms found in CD16^+^ monocytes is depicted in the box at the top. **(D)** Heat map of immune system-associated genes in CD16^+^ monocytes predicted by the GO-shuffling approach. The mean gene expression per donor is represented as a z-transformed value (across all donors). Columns and rows of the heat map are sorted by hierarchical clustering. **(E)** Violin plots of the expression of genes, found as DE in AMs, in blood monocytes based on scRNA-seq data. The plots show the expression across the donors, whereby the donors were downsampled to the same number of cells, followed by downsampling to the same number of cells between COPD and control. The plots display cells with an expression > 0. BALF = bronchoalveolar lavage fluid; AM = alveolar macrophage; mono = monocyte; MΦ = macrophage; proliferat. = proliferating; intermed. = intermediate; Pat. = patient; Ctrl. = control.

## Supplementary Figure legends

**Table S1: Patient overview with clinical parameters**

**Table S2: Overview of used FACS panels**

**Table S3: Cell numbers for the blood and BALF scRNA-seq dataset of the integrated analysis (related to Figure 2 – 7)**

## Material and Methods

### Human Specimens

Human studies were approved by the ethics committees of the University of Bonn and University hospital Bonn (local ethics vote 076/16). All patients provided written informed consent according to the Declaration of Helsinki before specimens were collected. Each individual included in this study was diagnosed and the disease stage was stratified according to the recommendations of the global initiative for chronic obstructive lung disease (GOLD) (COPD recommendations, 2020), with a ratio of post-bronchodilator (salbutamol 400 µg) forced expiratory volume in 1 s (FEV1) to forced vital capacity (FVC) of less than 0.7, and moderate airflow limitation (50% <= FEV1 < 80%). For scRNA-seq, the eligible patients were aged 40 years or older and were either current or ex-smokers. We focused on GOLD 2 patients (**Table S1**) and used stringent exclusion criteria for patients primarily diagnosed with asthma, clinically significant cardiovascular disorders or laboratory abnormalities and unstable concurrent disease (e.g. exacerbation of disease) that could have affected safety (as judged by the investigator). Individuals suffering from chronic cough without any signs of severe lung pathophysiology or subnormal lung functions served as control donors.

### Isolation of cells from BALF

Human BALF was obtained from patients with or without COPD via bronchoscopy (at the University hospital Bonn). BALF samples were once washed with PBS supplemented with 1 mM EDTA followed by washing with PBS supplemented with 2% fetal calf serum (FCS) and 1 mM EDTA. Throughout the isolation process, the samples were kept at 4°C and centrifugation steps performed at 300 g for 10 min. To exclude any macroscopic non-cellular particles and non-immune cells from further analyses, immune cells were enriched with MACS columns (Miltenyi) by using CD45 microbeads according to manufacturer’s instructions.

### Isolation of PBMCs and granulocytes

For the assessment of relationship analysis of the myeloid cell compartment in BALF with cells from the systemic circulation, we obtained venipuncture blood on the day of bronchoscopy. PBMC were obtained by Pancoll density centrifugation. After harvesting PBMC from the interphase, all further steps were conducted at 4°C. Granulocytes were recovered from the granulocyte/erythrocyte fraction using cold ACK (ammonium chloride potassium) lysing buffer to lyse erythrocytes, followed by a washing step with PBS supplemented with 2% FCS and 1 mM EDTA. To assess the granulocyte fraction in further analyses, it was mixed with the PBMC fraction in the ratio PBMC:granulocytes = 2:1.

### Flow cytometric data generation

Cells were resuspended in PBS supplemented with 2% FCS and 1 mM EDTA for surface marker staining (**Table S2**). To distinguish live from dead cells, the cells were incubated with LIVE/DEAD Fixable Dead Cell Stain Kit (1:1000). After washing, human FcR blocking reagent was included to reduce unspecific staining. Next, surface antibodies were added and cells were washed and analyzed either on BD FACSAria III (Becton Dickinson) for acquisition and sorting or on BD FACSCanto II (Becton Dickinson) for acquisition only (**Table S2**).

### Flow cytometric data analysis

Preliminary data analysis was performed using FlowJo software (version 10). The package ‘flowCore’ (version 1.46.2, Ellis et al., 2019) was used to import the compensated data into R. For dimensionality reduction with UMAP implementation in R (version 0.2.1.0, McInnes et al., 2018), fluorescence parameters were transformed with logicleTransform (Becht et al., 2019; Parks et al., 2006). Subsequent clustering of the dataset was performed with the PhenoGraph algorithm implemented in the ‘Rphenograph’ package (version 0.99.1, Levine et al., 2015). Based on marker detection, the identified clusters were assigned to cell-types. To unify and simplify the analysis across multiple datasets, an annotated dataset was defined as the reference and the other flow cytometry datasets were projected onto its UMAP coordinates.

### MitoStress assay on Seahorse

For the analysis of the metabolic state of donor-derived macrophages, AMs were isolated by adherence (with prior depletion of granulocytes using CD66b microbeads (Miltenyi)) and loaded onto the Seahorse XFe96 Analyzer (Agilent). After 3 cycles of baseline measurement, the cells were subsequently injected with Oligomycin (1:1000), FCCP (1:500) and finally a combination of Antimycin A and Rotenone (both 1:2000). Following each injection, oxygen consumption rate (OCR) was measured for 3 cycles.

### Migration Assay

Migration was analyzed in 24 well transwell plate containing a 8 µm polycarbonate membrane. FACS sorted AMs were suspended in 300 µL starvation medium (RPMI 1640 medium supplemented with 0.5% FCS and 1% penicillin/streptomycin) and 50,000 AMs were seeded in each upper well, while the lower chamber was filled with 700 µL starvation medium only. After an incubation of 1 h in a 37°C incubator, the medium in the upper chamber was exchanged with 300 µL fresh starvation medium and the medium in the lower chamber with 700 µL starvation medium supplemented with 100 ng/mL recombinant human CCL3. The seeded AMs were incubated at 37°C overnight. Next, cells on the upper filter surface were removed with a cotton swab. Transmigrated cells on lower filter surface were incubated with 2 µM CFSE and imaging of cells was performed using an inverted fluorescent microscope (Nikon).

### Measurement of proteins in BALF

After isolation of cells (see above), the supernatant of BALF samples of both COPD patients and controls were collected and frozen at −80 °C before proteomics measurement. Protein levels from cell-free BALF samples were determined using the INFLAMMATION panel from Olink Proteomics.

### Lipidomics of macrophages in BALF

AMs were sorted, washed with PBS and with 150 mM ammonium acetate in a glass tube, pelleted (300 g with slow brake), and frozen at −80°C until analysis. To the pellet, 500 µL of extraction mix (CHCl_3_/MeOH 1/5 containing internal standards: 210 pmol PE(31:1), 396 pmol PC(31:1), 98 pmol PS(31:1), 84 pmol PI(34:0), 56 pmol PA(31:1), 51 pmol PG (28:0), 28 pmol CL(56:0), 39 pmol LPA (17:0), 35 pmol LPC(17:1), 38 pmol LPE (17:1), 32 pmol Cer(17:0), 99 pmol SM(17:0),55 pmol GlcCer(12:0), 14 pmol GM3 (18:0-D3), 359 pmol TG(47:1), 111 pmol CE(17:1), 64 pmol DG(31:1), 103 pmol MG(17:1), 724 pmol Chol(d6), 45 pmol Car(15:0)) were added and each sample sonicated for 2 min followed by centrifugation at 20,000 g for 2 min. The supernatant was collected into a new tube and 200 µL chloroform and 800 µL 1% AcOH in H_2_O were added. The sample was then briefly shaken and spun for 2 min at 20,000 g for 2 min. 200 µL chloroform and 800 µL 1% AcOH in H_2_O were added to the supernatant, briefly shaken and spun for 2 min at 20,000 g. The lower phase was transferred into a new tube and evaporated in a speed vac (45°C, 10 min). Spray buffer (500 µL of 8/5/1 2-propanol/MeOH/H_2_O, 10 mM ammonium acetate) was added, sonicated for 5 min and infused at 10 µL/min into a Thermo Q Exactive Plus spectrometer (Thermo Fisher Scientific) equipped with the HESI II ion source for shotgun lipidomics. MS1 spectra (res. 280000) were recorded in 100 m/z windows from 200 – 1200 m/z (pos.) and 200 – 1700 m/z (neg.) followed by recording MS/MS spectra (res. 70000) by data independent acquisition in 1 m/z windows from 200 – 1200 (pos.) and 200 – 1700 (neg.) m/z.

### Nanodroplet-based scRNA-seq

For each donor, 10,000 BALF or blood-derived cells were loaded onto the Chromium™ Controller instrument (10x Genomics) using the Chromium™ Single Cell A Chip Kit together with the Chromium™ Gel Bead Kit v2 following the manufacturer’s recommendations. Libraries were prepared using Chromium™ Single Cell 3’ Library Kit v2 according to manufacturer’s recommendations and sequenced paired-end as followed: Read 1 26 cycles, i7 index 8 cycles and Read 2 56 cycles on a NextSeq500 instrument (Illumina) using High Output v2.1 chemistry. Single-cell data was demultiplexed and converted into fastq format using bcl2fastq2 (v2.20).

### Generation of scRNA-seq libraries using Seq-Well

Seq-Well libraries were generated as described by Gierahn *et al*. (Gierahn et al., 2017), with 20,000 BALF or blood cells per donor loaded on an array. To reduce library costs, we produced homemade Tn5 transposase according to Picelli *et al*. (Picelli et al., 2014). Seq-Well libraries were equimolarly pooled and clustered at 1.4pM concentration with 10% PhiX using High Output v2.1 chemistry on a NextSeq500 system. Sequencing was performed paired-end as followed: custom Drop-Seq Read 1 primer for 21 cycles, 8 cycles for the i7 index and 61 cycles for Read 2. Single-cell data were demultiplexed using bcl2fastq2 (v2.20).

### Processing of scRNA-seq raw data

For preprocessing, the generated fastq files from both Chromium™ and Seq-Well were loaded into a data pre-processing pipeline (version 0.31, available at https://github.com/Hoohm/dropSeqPipe) that relies on Drop-seq tools provided by the McCarroll lab (Macosko et al., 2015). STAR alignment within the pipeline was performed using the human GENCODE reference genome and transcriptome hg38 release 27 (Harrow et al., 2012). The resulting datasets were then imported into the R package ‘Seurat’ (v.3.0.0, Butler et al., 2018) for downstream analyses.

### Quality control of scRNA-seq data

We defined cells and genes to be included for further analyses by the following criteria for each donor separately: (1) Only genes that were found in at least 3 cells were kept; (2) 100 expressed genes was used to keep cells for further analyses; (3) With regard to the rate of endogenous-to-mitochondrial counts per cell, blood cells with a rate > 5% and lavage cells with a rate >10% were excluded. For the comparison of scRNA-seq methods for clinical applications, these quality control filters resulted in a Chromium™ dataset of 13,909 cells (BALF = 7,960 cells; blood = 5,949 cells) across 22,701 genes and a Seq-Well dataset comprised of 34,622 cells (BALF = 20,106 cells; blood = 14,516 cells) across 21,644 genes. For the integrated analysis of Seq-Well data from COPD GOLD 2 patients and control donors, we obtained a Seq-Well dataset of 60,925 lavage cells across 25,348 genes and 54,569 blood cells across 23,056 genes (**Table S3**).

### Dataset integration and dimensionality reduction of scRNA-seq data

Cells were log-normalized using the CPM-normalization with a scale factor of 10,000. Next, the genes with the highest cell-to-cell variability in the dataset were determined by calculating the top 2,000 most variable genes by selecting the ‘vst’ method of the ‘FindVariableFeatures’ function in Seurat. We used data integration based on ‘anchors’ across batches (Stuart et al., 2019) to harmonize and integrate the different datasets by using the Seurat implementation with the default settings (defined batch in the present dataset = donors). After scaling, the dimensionality of the data was reduced to 30 principal components (PCs) that was used as input for UMAP representation.

### Cell-type annotation based on reference transcriptomic datasets

For the comparison of the datasets generated by the two different scRNA-seq technologies (see above), we developed a slightly modified Python implementation of SingleR (Aran et al., 2019b). As a reference for SingleR, we used data from both Blueprint+ENCODE (Dunham et al., 2012; Stunnenberg et al., 2016) and the Human Primary Cell Atlas (HPCA) (Mabbott et al., 2013).

As another cell annotation approach, we developed the tool GenExPro (Gene Expression Profiler) that will be published elsewhere in detail. The basic idea of GenExPro is to compare the mean vector of gene expressions from a cluster of cells in the single-cell dataset to the expression profiles of a reference dataset of expression profiles with annotated cell types. To this end, the GenExPro method fits a multiple linear regression for each defined expression vector. The covariates in this regression are the reference expression vectors for each cell type that were obtained from the CIBERSORT algorithm (Newman et al., 2015).

### Consolidation of cell-type annotation using machine learning

To aggregate and consolidate the initial cell-type annotation, we trained a Gradient Boosting Classifier on the combined data of all datasets to classify each cell into a cell type. We used an implementation of the Gradient Boosting algorithm from scikit-learn (version 0.19.1, Pedregosa et al., 2012). Our aim was to apply the classifier to all cells in our data. However, as no distinct training data was available, we conducted a 3-fold cross-validation. In this procedure, two random thirds of a data set were used as training data, and the model assigned cell type names to the remaining cells.

### Clustering of the integrated scRNA-seq datasets

The cellular heterogeneity of the integrated datasets was determined using a shared nearest neighbor (SNN)-graph based clustering algorithm implemented in the Seurat pipeline. For both the BALF and the blood data, we used the first 30 principle components as input and set the resolution to 0.7 and 0.6, respectively. The default setting for number of neighbors were used (k=20).

### Marker gene identification of scRNA-seq data

Marker genes for identified cell types/clusters were calculated using a Wilcoxon rank sum test for differential gene expression implemented in Seurat. The significance threshold for marker genes were set to an adjusted p-value smaller than 0.001 and the logarithmic fold change cutoff to at least 0.4. In addition, the detected marker genes should have been expressed in at least 50% of the cells within the respective cell types/clusters.

### ‘Gene set distance’ analysis of annotated cell types (GO-shuffling)

Gene set annotations were downloaded from the Molecular Signatures Database v7.0 (MSigDB) and comprised gene sets from the Kyoto Encyclopedia of Genes and Genomes (KEGG) (Kanehisa, 2019) database, the Pathway Interaction Database (PID) (Schaefer et al., 2009), the Reactome Pathway database (Fabregat et al., 2018), Hallmark gene sets (Liberzon et al., 2015), BioCarta Pathways (Nishimura, 2001) and Gene Ontology (GO) (Ashburner et al., 2000; Carbon et al., 2019). In addition, we retrieved gene sets from WikiPathways (Slenter et al., 2018). Gene sets were taken into account that were present with a minimum of 3 genes. For each gene set, the Euclidean distance between all donors was calculated using the get_dist function from the R package ‘factoextra’ (version 1.0.5). The “gene set distance” was then defined as followed: 

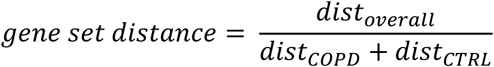

### Modeling of metabolic pathways based on scRNA-seq data

The metabolic landscape of AMs was modelled using the Compass method (version 0.9.5, Wagner et al., 2020; Wang et al., 2020) by leaving the standard settings unaltered. As input, we simplified the single-cell data of the AMs by using the ‘applyMicroClustering’ function of the R package ‘VISION’ (version 2.1.0, DeTomaso et al., 2019), resulting in approximately 20 microclusters per patient. Next, we applied Compass to the microclusters for each donor separately. The output tables were concatenated and transformed as described in the Compass manuscript, except for disabling the division into meta-reactions. To determine which reactions and metabolites are significantly different between control donors and COPD patients, we performed Wilcoxon rank sum tests on Compass scores.

### Cell cycle state analysis of scRNA-Seq data

To categorize the cells within the AM clusters into the respective cell cycle states, we applied the ‘CellCycleScoring’ function of Seurat and substantiated the results using the ‘cyclone’ function (Scialdone et al., 2015) implemented in the R package ‘scran’ (version 1.10.2, Lun et al., 2016).

### Gene set variation analysis

To predict the functions of the AM states, we performed gene set variation analysis (GSVA) (Hänzelmann et al., 2013b) by using the R package ‘GSVA’ (version 1.30.0). For the GSVA input expression table, we calculated the sum of the expression of normalized scRNA-seq data for each patient in any AM states. As gene sets we used the gene set collection described in the section ‘GO-shuffling’ and additionally included the ‘ImmuneSigDB’ collection of MsigDB. The GSVA results per donor were combined for the respective AM state using a Borda rank and the top 250 ranked gene sets per subtype were visualized in an UpSet plot using the R package ‘UpSetR’ (version 1.3.3, Conway et al., 2017)).

### AUCell for gene set enrichment analysis

Enrichment of gene sets was performed using the ‘AUCell’ method (Aibar et al., 2017) implemented in the package (version 1.4.1) in R. We set the threshold for the calculation of the area under the curve (AUC) to the top 3% of the ranked genes and normalized the maximum possible AUC to 1. The resulting AUC values were subsequently visualized in a violin plot.

### Distribution-free differential expression (DE) analysis across patient groups

To analyze the differences between patient and control cohort we developed a distribution-free test, which preserves patient and cell information. In contrast to available methods, it avoids the use of mini-bulk, the pooling of cells from different patients, and distribution assumptions. As input, we use the afore-computed cell cluster information and the normalized single-cell data.

For each AM cluster a differential expression between patient and control cohort was performed. Therefore, individuals not possessing cells in a cluster – which happened in a few cases – and genes expressed in less than 10% of cells were disregarded for the analysis of this cluster. For each gene, the differences between all possible pairs of patients and controls was assessed using the nonparametric Wilcoxon rank sum test. To assess the differences between patient groups, the median Wilcoxon score of the pair-wise tests was used considered as test statistic.

To assess if the observed value of the test statistic was significant, the probability of observing an equally or more extreme value of the test statistic under the null hypothesis was evaluated. The null hypothesis was that there is no difference between the two groups. The exact null distribution was evaluated with the permutation test, taking all possible permutations into account. For all possible permutations the afore-described test statistic – the median Wilcoxon score – was evaluated. The distribution of the test statistic over all permutations provided the null distribution, since reshuffling of patients should not be significant under the null hypothesis. The p-value for the observed group assignment was then the fraction of permutations that led to an equal or more extreme value of the test statistic than the value of the test statistic of the observed patient arrangement.

### Gene set enrichment analysis

GSEA was performed to identify shared common biological functions by groups of DE genes. The web-tool ‘g:Profiler’ (Reimand et al., 2007) was used to perform the functional profiling of the DE genes. As multiple-testing correction method, g:Profiler’s in-house g:SCS algorithm was chosen.

### Cell-to-cell communication

Potential cell-cell-interactions were inferred using ‘CellPhoneDB’ (version 2.1.1, Efremova et al., 2019; Vento-Tormo et al., 2018). As input, we used the normalized gene expression matrix of control and COPD patients that was filtered separately for cell types that contained ≥ 10 cells per patient. Genes were filtered for being expressed in ≥ 5% of a respective cell type. To run CellPhoneDB, the following parameters were set: --iterations=1,000 --pvalue=0.1 --result-precision=10.

For ‘NicheNet’ analysis (version 0.1.0, Bonnardel et al., 2019; Browaeys et al., 2019). we accepted all genes that were expressed in >5% of any cell type within the COPD group and which matched at least one receptor from the genes expressed in > 5% of the mono-like macrophages in the COPD group. As input genes to infer the ligand activity score from, we defined all DE genes with a median Wilcoxon score < (−0.75) and p value of the median Wilcox score <0.05. As background genes, we defined all genes that are not DE in mono-like macrophages and expressed in > 5% of mono-like macrophages. For ligand prioritization, we selected the top 20 genes with the highest PCC or AUPR resulting in 26 top ligands. To not miss out any cell type-specific cell-cell-interaction, we additionally used every cell type separately as sender cell and chose the top 7 genes according to the PCC and added these to the top ligands resulting in 32 top genes.

### Monocyte-to-macrophage trajectory analysis

To generate a joint embedding of BAL and blood samples, the data were jointly pre-processed using ‘Scanpy’ (version 1.4.3, Wolf et al., 2018) on AnnData (version 0.6.22). In concordance with previous analysis, cells from BALF were filtered out if the fraction of mitochondrial reads exceeded 0.1, and a threshold of 0.05 was used for blood samples. Genes that were expressed in fewer than 200 cells were also filtered out. Following previously published best-practices (Luecken and Theis, 2019) we used scran normalization via the computeSumFactors function on the joint object. Spliced and unspliced counts were mapped to this object using scVelo (version 0.1.24 commit e45a65a, Bergen et al., 2019). Quality control for spliced and unspliced counts was performed by removing cells with fewer than 20 spliced and/or 10 unspliced counts. Subsequent normalization by total counts and log-transformation was performed via the filter_and_normalize function from scVelo.

The joint embedding of BAL and blood cells was generated by taking the top 4000 highly variable genes (HVGs) that were shared by most batches. This was done using the hvg_batch function from the single-cell data integration benchmarking package ‘scIB’ (published separately). This function computes the top 4000 HVGs per batch (here: donor) using Scanpy’s highly_variable_genes function with method cell_ranger and ranks these by the number of batches these genes are highly variable in, and by their mean dispersion over all batches. In this list, the top 4000 genes are selected. On this gene set, we computed the top 50 principle components and used Euclidean distance on these to compute a kNN graph with a k=15. ‘UMAP’ (version 0.3.9, McInnes et al., 2018) was used to visualize the results.

Due to an observed batch effect when performing RNA velocity analysis across patients, we ran scVelo per patient and aggregated the individual patient velocities to create a joint velocity embedding. For each donor spliced and unspliced counts were smoothed using the moments function, velocity genes were selected by a stringent log likelihood threshold of 0.1 (between 45 and 172 genes per donor), and the dynamical scVelo model was fit. The resulting inferred single-cell velocities were projected onto the joint UMAP computed from all donors by running velocity_graph on the concatenated object.

Furthermore, partition-based graph abstraction (‘PAGA’, Wolf et al., 2019) was used to assess the connectivity of cell identity clusters that were suggested to show transitions by RNA velocity. To robustly assess the connectivity of cell identity clusters across donors, we performed PAGA analysis per donor. We computed a kNN graph with Scanpy’s neighbors function (k=15) per donor using the joint PCA embedding across donors and ran the paga function on this graph. We used the resulting PAGA connectivities as a statistical test of kNN-graph connectivity between clusters. The median of PAGA connectivities over all donors with both blood and BAL samples was used as a PAGA distance metric.

### Statistical analysis

If not otherwise stated, the statistical evaluation was carried out in relation to the total sample size *n*. A t-test was used for *n* ≤ 10, otherwise a Wilcoxon rank-sum test was used.

